# Endogenous pioneer neutrophils release NETs during the swarming response in zebrafish

**DOI:** 10.1101/521450

**Authors:** Hannah M. Isles, Catherine A. Loynes, Noémie Hamilton, Clare F. Muir, Anastasia Kadochnikova, Katherine M. Henry, Visakan Kadirkamanathan, Stephen A. Renshaw, Philip M. Elks

## Abstract

Neutrophils are rapidly recruited to inflammatory sites where they coordinate their migration to form clusters, a process termed neutrophil swarming. The factors which modulate neutrophil swarming during its early stages are not fully understood, requiring the development of new *in vivo* models. Using transgenic zebrafish larvae to study endogenous neutrophil migration in a tissue damage model, we demonstrate that neutrophil swarming is a conserved process in zebrafish immunity, sharing essential features with mammalian systems. We show that neutrophil swarms initially develop around a pioneer neutrophil, in a three-phase sequence of events. By adopting a high-resolution confocal microscopy approach, we observed the release of cell fragments by early swarming neutrophils. We developed a neutrophil specific histone H2A transgenic reporter line *TgBAC(mpx:GFP)i114;Tg(lyz:H2A-mCherry)sh530* to study neutrophil extracellular traps (NETs), and found that endogenous neutrophils recruited to sites of tissue damage released NETs at the start of the swarming process. The optical transparency achieved using the zebrafish model has provided some of the highest resolution imaging of NET release *in vivo* to date. Using a combination of transgenic reporter lines and DNA intercalating agents, we demonstrate that pioneer neutrophils release extracellular traps during the swarming response, suggesting that cell death signalling via NETosis might be important in driving the swarming response.

## Introduction

A robust inflammatory response against invading pathogens or endogenous danger signals requires the coordination of multiple cellular and humoral components. Neutrophils are one of the first responders to tissue inflammation and rapidly home to inflamed tissue within hours of injury. Within inflamed tissue, neutrophils destroy pathogens ^1^ and clear wound debris ^2^, ultimately leading to the restoration of tissue homeostasis. The anti-microbial repertoire of neutrophils can cause substantial secondary tissue damage, thus neutrophil recruitment to, and removal from, inflammatory sites must be tightly controlled.

Neutrophils are recruited to sites of inflammation through a series of well-defined molecular events ^3–5^. During their recruitment, neutrophils are primed by pro-inflammatory signals including growth factors, inflammatory cytokines and chemoattractants. Neutrophils are capable of integrating host- and pathogen-derived environmental signals, resulting in their polarisation and migration towards the initiating inflammatory stimulus ^6^. Within the interstitium, neutrophils coordinate their migration patterns to form clusters in several models of sterile-inflammation and infection ^7–13^. The parallels between these cellular behaviours and migration patterns seen in insects has led to use of the term “swarming”.

A series of sequential phases leading to neutrophil swarming have been described in murine models; the initial migration of ‘pioneer’ or neutrophils proximal to the wound site (scouting) is followed by large scale synchronised migration of neutrophils from distant regions (amplification) leading to neutrophil clustering (stabilisation) and eventual resolution ^7–10^. The initial arrest and death of early recruited pioneer neutrophils correlates with the onset of neutrophil swarming ^8,9,13^, which is mediated by many chemoattractants including lipid and protein mediators, with a dominant role for the lipid leukotriene B4 (LTB4) identified *in vivo* ^7,8^. It is likely that swarming neutrophils respond to an amplified signal initiated by pioneer neutrophils, however the precise nature of these signals and mode of cell death which pioneer neutrophils undergo remains to be determined.

At sites of inflammation neutrophil death can be modulated by extracellular stimuli such as proinflammatory cytokines, pathogens, toxic tissue constituents, and platelets ^14^. Depending on the inducing stimulus, neutrophils ultimately die through different modes of cell death which manifest with unique macroscopic morphological changes ^15^. Controlled cell death including autophagy and apoptosis involve a clean and non-inflammatory mode of cellular destruction, whilst accidental cell death such as necrosis results in the spilling of cellular contents and release of pro-inflammatory signals ^16^. Neutrophils can also die following their production of extracellular traps (NETosis) composed of DNA and histones embedded with granular and cytoplasmic proteins, which are able to capture and kill pathogens extracellularly ^17^. Neutrophils release NETs following a series of intracellular changes resulting in chromatin decondensation, breakdown of the nuclear envelope and mixing of DNA with granular and cytoplasmic proteins ^18^. The uptake of DNA intercalating agent propidium iodide and the expulsion of vesicles from pioneer neutrophils precedes the onset of swarming in murine models ^13^, suggesting that accidental cell death by necrosis or extracellular DNA release during NETosis are likely pioneer cell death mechanisms and represent interesting avenues for exploration.

The zebrafish (*Danio rerio*) is a powerful model organism in which to study neutrophil function that has been used extensively to study neutrophil migration towards and away from sites of sterile inflammation ^19–21^. The optical transparency of zebrafish embryos allows for the tracking of endogenous neutrophils at wound sites within minutes following injury in transgenic reporter lines ^22^. In this study, we use zebrafish larvae to study pioneer neutrophil behaviour prior to the onset of swarming. We use both inflammation and infection assays to demonstrate that neutrophil swarming is conserved in zebrafish immunity, indicating importance of this neutrophil behaviour across evolution. We define a three-stage sequence of migration events which leads to the swarming of endogenous neutrophils within the inflamed tissue and verify that the neutrophil relay signal, LTB4, is required for amplification of neutrophil recruitment. Importantly, we show that a single pioneer neutrophil is sufficient to induce a swarming response and that this neutrophil adopts a rounded, non-motile morphology distinct from other neutrophils within the inflamed tissue. We develop a transgenic reporter for NETs and perform live imaging of neutrophil extracellular trap release. We study pioneer neutrophil cell death using cell viability assays and transgenic reporters for cell death, and identify that pioneer neutrophils are viable prior to the onset of swarming, but die following the release extracellular traps from within the swarm centre.

## Results

### Neutrophil swarming is conserved in zebrafish immunity

Neutrophil swarming is characterised by the highly directed and coordinated movement of neutrophils to sites of infection or injury followed by accumulation and clustering ^23^. To determine whether neutrophil swarming is conserved in zebrafish immunity neutrophil mobilisation to both inflammatory and infectious stimuli was studied. Neutrophil responses to inflammatory stimuli were assessed by tail fin transection of 3 days post fertilisation (dpf) *mpx*:GFP larvae, and tracking neutrophil migration using fluorescence microscopy during recruitment (0-6 hours post injury, Supplementary Figure 1). Analysis of migration patterns of neutrophils recruited to the wound site identified three outcomes: 1) persistent neutrophil swarming reminiscent of neutrophil swarming reported in mammalian systems ^7–10^ (Figure 1A-C, Supplemental Movie 1); 2) shorter lived transient neutrophil swarms which dissipated and reformed multiple times within the imaging period (Supplemental Figure 2, Supplemental Movie 2); 3) no coordinated migration and no swarm formation (Supplemental Movie 3). Persistent swarming was defined as the formation of clusters which grew by the coordinated migration of individual neutrophils (Figure 1C). Persistent swarms were observed from 40 minutes post injury (Supplemental Figure 3A) and remained stable for, on average, 2.17 hours ± 0.32 (Supplemental Figure 3B). Persistent neutrophil swarms were observed in 50% of larvae, transient swarms (persisting for <1 hour) were seen in 14% of larvae, and 36% of larvae showed no evidence of swarming behaviour within the imaging period (Figure 1D).

**Figure 1.**
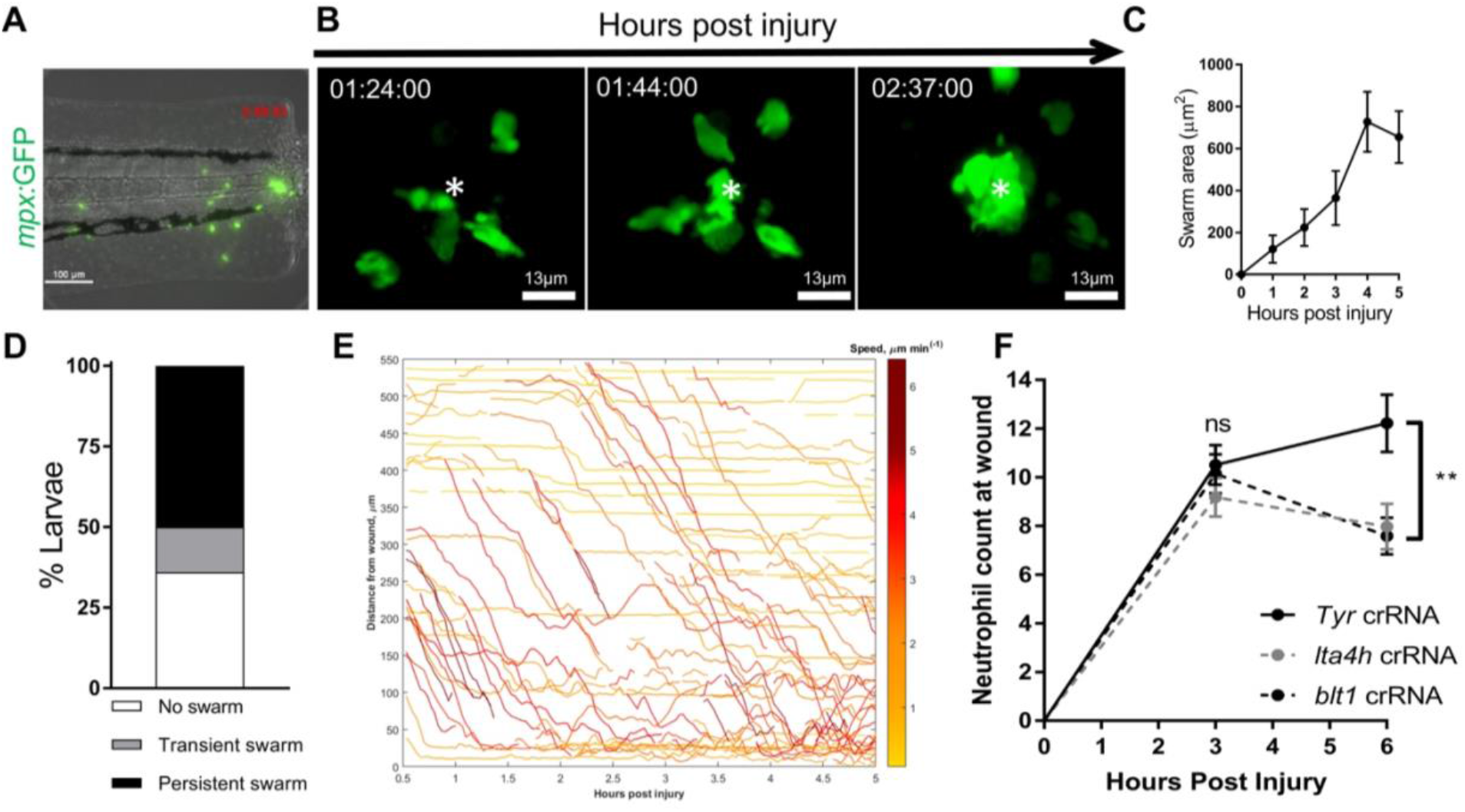
Neutrophil swarming is conserved in the zebrafish tissue damage response. **A-C** Zebrafish neutrophils swarm at sites of tissue damage. **A** Representative image illustrating neutrophils swarming at the wound site following tail fin transection in 3dpf *mpx*:GFP larvae. Image was taken using 20x magnification on a TE2000U inverted microscope (Nikon). Time stamp shown is relative to the start of the imaging period at 30 minutes post injury and is h:mm:ss. **B** 3D reconstruction time course illustrating neutrophils swarming at the wound site (swarm centre is highlighted by white asterix). Imaging was performed using a 40X objective spinning disk confocal microscope (Perkin Elmer). Time stamps shown are relative to time post injury and are in hh:mm:ss. **C.** Area of neutrophil swarms measured at hourly intervals during the 5 hour imaging period. Error bars shown are mean ± SEM, n=7. **D** Frequency of neutrophil swarming behaviour observed at the wound site within 5 hours following injury, n=14. **E-F** Relay signalling through LTB4 is required for neutrophil recruitment. **E** Distance time plot demonstrating the early recruitment of neutrophils proximal to the wound site (<350μm) followed by the later recruitment of more distant neutrophils. Tracks are colour coded based on their average speed (μm/minute). F CRISPR/Cas9-mediated knockdown of LTB4 signalling reduces late neutrophil recruitment. Neutrophil counts at the wound site in control *tyr* crRNA injected larvae (black line), *lta4h* crRNA injected larave (grey dotted line), and *blt1* crRNA injected larvae (black dotted line) at 3 and 6hpi. Error bars shown are mean ± SEM. Groups were analysed using an ordinary one-way ANOVA and adjusted using Tukeys multi comparison test. **p>0.008 n=45 from 3 independent repeats.

In mammalian neutrophil swarming, biphasic neutrophil responses are modulated by the lipid LTB4 ^8^. During the imaging period, two waves of neutrophil recruitment were observed: the early migration of neutrophils proximal to the wound site between 0.5-2hpi, followed by a later influx of neutrophils from more distant sites (Figure 1E). We investigated the requirement for LTB4 in neutrophil chemotaxis towards the wound site in zebrafish using the CRISPR/Cas9 system. Biosynthesis of LTB4 in zebrafish occurs through fatty acid metabolism of arachidonic acid via common intermediates, resulting in the production of LTB4 by the enzyme leukotriene A4 hydrolase (Lta4h), encoded by the gene *lta4h* ^24,25^. Zebrafish have three LTB4 receptors; the high affinity Blt1 receptor and two low affinity receptors Blt2a and Blt2b, of which zebrafish neutrophils predominantly express *blt1* (Supplemental Figure 4A-B). Using Cas9 protein with guide RNAs (crRNAs) to target the *lta4h* and *blt1* genes, early neutrophil recruitment (3hpi) and late neutrophil responses (6hpi) to tail fin injury were assessed. Early neutrophil recruitment to the wound site at 3hpi was similar between control (*tyrosinase*) ^26,27^, *blt1* and *lta4h* crRNA injected larvae (Figure 1F), suggesting that LTB4 signalling is not required for early neutrophil responses. At 6hpi, neutrophil recruitment in *blt1* and *lta4h* crRNA injected larvae was significantly lower than control (*tyr*) crRNA injected larvae (Figure 1F). These results are in agreement with data from mouse ^8^ and human neutrophils ^7^, supporting a requirement for LTB4 signalling in neutrophil recruitment at the later stages.

After determining that swarming was a conserved component of the tissue damage response in zebrafish, neutrophil responses to infectious stimuli were assessed. Otic vesicle injection of *Staphylococcus aureus*, a gram positive bacteria which induces a robust neutrophil swarming response in mammalian neutrophils ^28^, induced robust neutrophil recruitment (21±2 neutrophils), which was not seen in larvae injected with a PBS control (1±0.3 neutrophils) (Figure 2A-B). Neutrophils within otic vesicles infected with *S. aureus*, but not PBS, coordinated their migration to form swarms in tissue regions containing bacteria, which at 6hpi had an average volume of 48.1mm^3^ (Figure 2C-D, Supplemental Movie 4). The identification of neutrophil swarming in response to inflammatory and infectious stimuli demonstrates that neutrophil swarming is a conserved component of zebrafish immunity. We therefore used the zebrafish model to understand how endogenous swarms are initiated *in vivo*.

**Figure 2.**
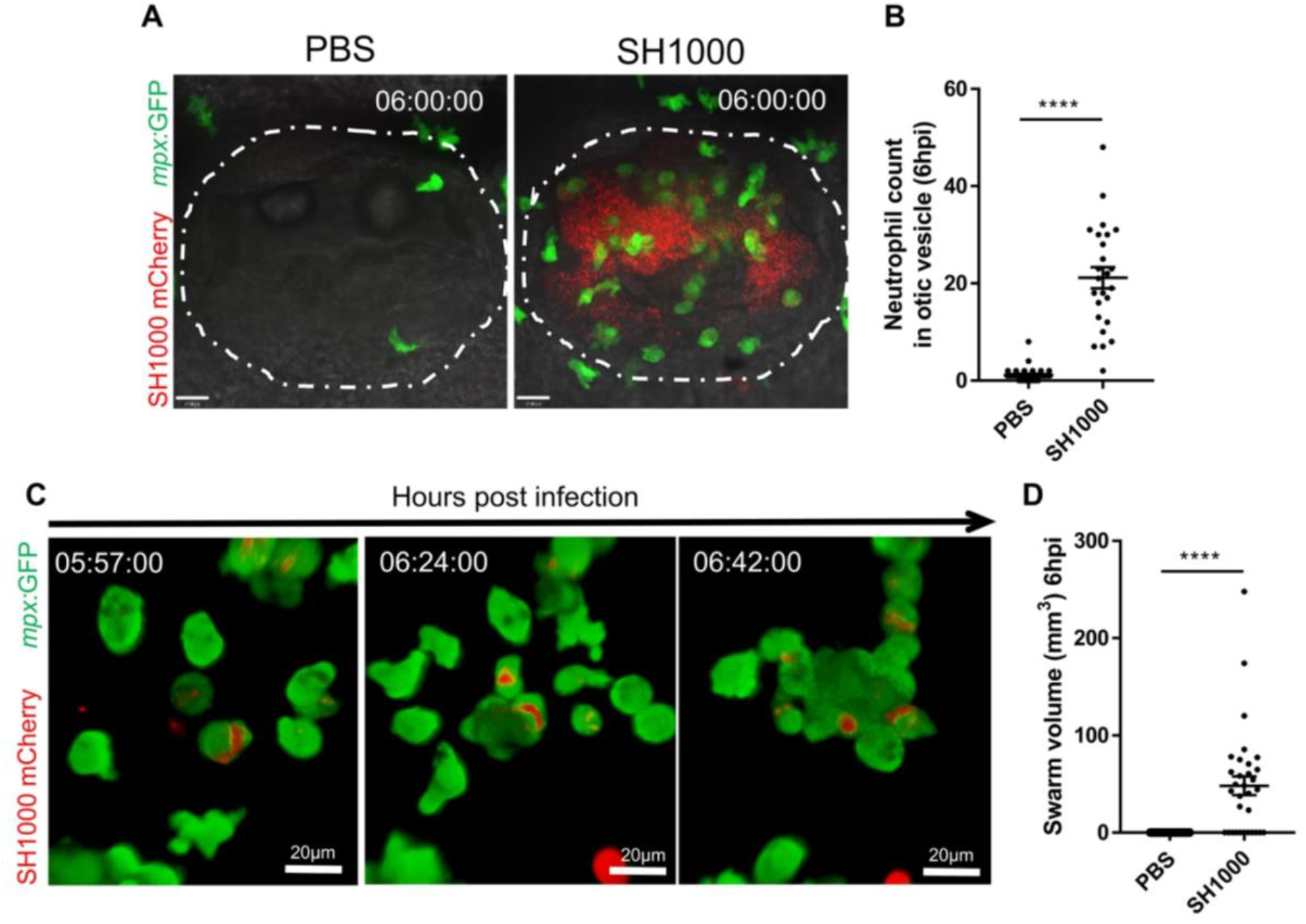
Zebrafish neutrophils swarm at *S. aureus* infection. **A-B** Neutrophils mobilised to the otic vesicle in response to *S. aureus*. Otic vesicles of 2dpf *mpx*:GFP larvae injected with a PBS vehicle control or 2500 cfu *S. aureus* SH1000 pMV158mCherry. **A** Representative image illustrating neutrophil recruitment (green) to otic vesicle infected with *S. aureus* (red). Otic vesicles are highlighted by white dashed area. Time stamps shown are hh:mm relative to time post infection. **B** Number of neutrophils mobilised to the otic vesicle at 6hpi. Error bars shown are mean ± SEM (****p>0.0001 from an un-paired t-test, n=32 from 3 independent repeats). **C** 3D reconstruction time course illustrating neutrophils swarming within the otic vesicle of 2dpf *mpx*:GFP larvae injected with 2500 cfu *S. aureus* SH1000 pMV158mCherry. Imaging was performed using a 20X objective spinning disk confocal microscope. Time stamps shown are hh:mm:ss relative to time post injection. **D** Volume of neutrophil swarms measured within otic vesicle at 6hpi. A volume of zero corresponds to no swarm observed. Error bars shown are mean ± SEM (****p>0.0001 from an un-paired t-test, n=32 from 3 independent repeats).

### Neutrophil swarms develop around a pioneer neutrophil in three sequential stages

Neutrophil swarms in mammalian models of inflammation grow by large-scale migration of neutrophils towards early recruited ‘pioneers’, which likely release additional signals to initiate the swarming response ^8–10^. To assess whether a similar process was relevant to endogenous neutrophils, we analysed the migration patterns of neutrophils in the time period leading to swarm formation, where the presence of one individual neutrophil was identified in the tissue region which became the swarm centre in 100% of swarming events examined (Figure 3A). Because a single neutrophil always initiated a swarm, we termed it the pioneer neutrophil by analogy to mammalian descriptions. In 100% of swarm initiation events examined, the pioneer neutrophil was the focal point of migration for early swarming neutrophils. Early swarming neutrophils directed their migration towards the pioneer neutrophil, whilst non-swarming neutrophils migrated with no apparent pattern within the wound region (Figure 3B-C, Supplemental Movie 5). Pioneer neutrophils were rounded and non-motile, which is illustrated by their higher circularity index and lower displacement compared to scouting neutrophils at the wound site in the same time period (Figure 4A-C). To investigate whether the round, non-motile morphology was distinct to pioneers, or common to all neutrophils upon arrival at the wound site, neutrophils migrating to the wound site were tracked during the scouting and the initiation phases. The speed, displacement and meandering index of pioneer neutrophils were significantly reduced in the initiation phase when compared to the scouting phase, whilst neutrophils migrating to the wound site within the same tissue region did not undergo this behavioural change (Figure 4D-F). These data demonstrate that pioneer neutrophils display a distinct morphology at the wound site prior to swarm formation, which is not seen in scouting neutrophils responding to chemoattractants produced at the wound edge.

**Figure 3.**
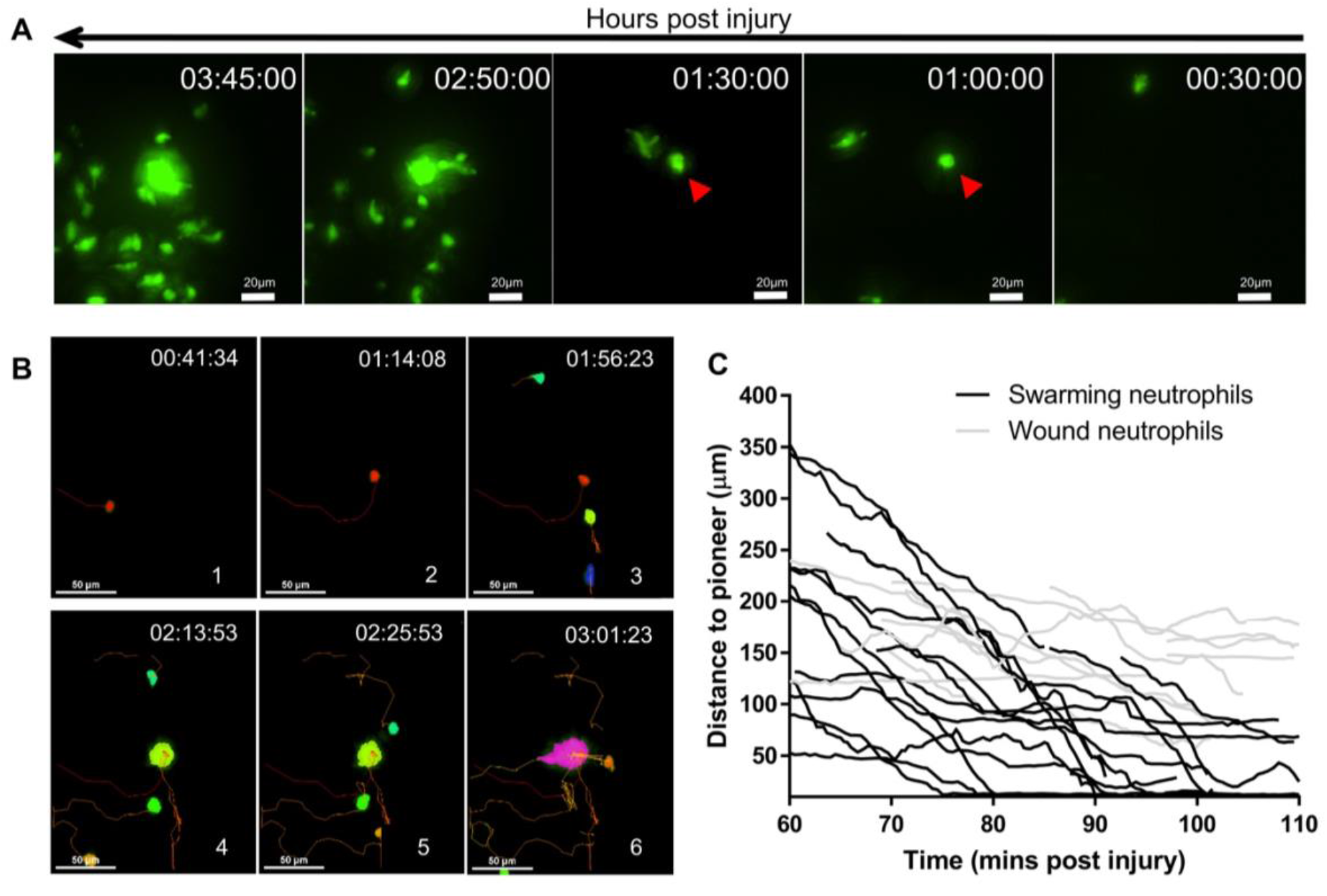
A pioneer neutrophil is the focal point of migration for swarming neutrophils. **A.** Reverse chronological time lapse sequence of a persistent neutrophil swarm where one individual neutrophil is visible in the swarm centre prior to neutrophil clustering (red arrows). Time stamps shown are hh:mm:ss relative to injury time. **B.** Chronological time lapse sequence of swarming neutrophil tracks. The migration of a pioneer neutrophil (red) to the wound site is observed (frames 1-2) followed by the directed migration of swarming neutrophils towards the pioneer, which is the focal point for migration (frames 3-5). The result of migration is the aggregation of neutrophils to form large clusters (frame 6). Tracks are coloured by time where red corresponds to early and yellow corresponds to late arriving neutrophils. **C.** Distance-time plot (DTP) of individual cell migration paths of swarming neutrophils (black tracks) and wound neutrophils at the wound site in the same time period (grey tracks). Tracks are relative to pioneer neutrophil position; swarming neutrophils migrate to the pioneer neutrophil whilst non-swarming neutrophils do not (n= 4 experimental repeats).

**Figure 4.**
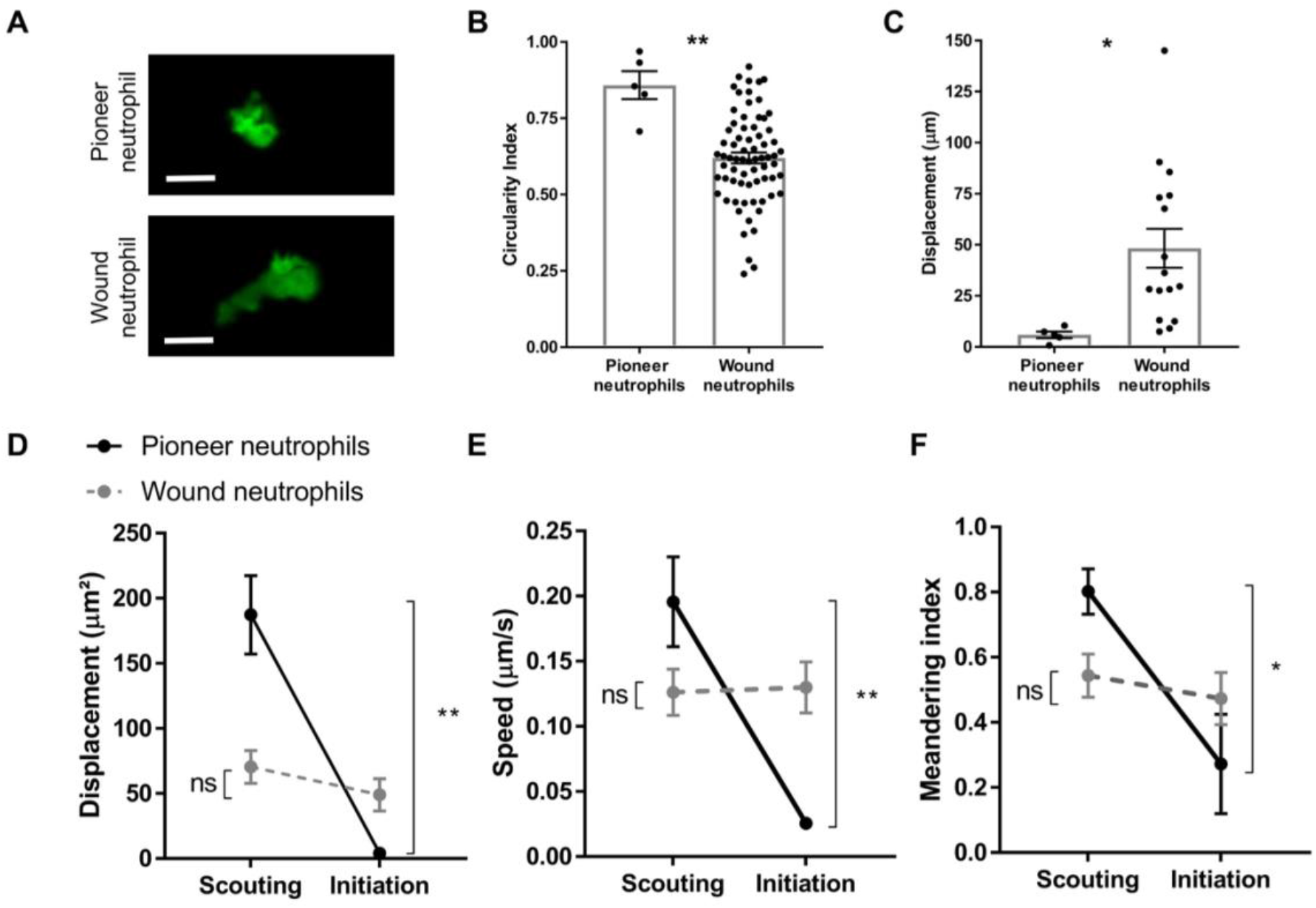
Pioneer neutrophils adopt a distinct rounded, non-motile morphology at the wound site. **A** Representative image of pioneer and non-pioneer neutrophil morphology. Images were taken using a 40X objective lens on a spinning disk confocal microscope (Perkin Elmer). Scale bars are 20 μm. **B-C** Quantification of pioneer neutrophil migration pattern in the frames preceding swarming.The circularity index (roundness) and displacement (movement) of pioneer neutrophils and wound neutrophils migrating at the wound site in the same time period (n=5, unpaired t-test where * p<0.05 and ** p<0.01). **D-F** Neutrophils were tracked from 30 minutes post injury. Parameters to study the migration patterns of pioneer and wound neutrophils were compared in the scouting and initiation phases. (D) Neutrophil displacement (the linear distance each neutrophil travelled). (E) Neutrophil speed. (F) Neutrophil meandering index (the displacement divided by the total length of the neutrophil track). (Error bars are mean ± SEM. Groups were analysed using a paired t-test *p<0.05 **p<0.01, n=5 independent repeats).

A series of sequential phases leading to neutrophil swarming has been described in mammalian systems ^8,9^, so we next determined the stages leading to swarming in zebrafish. Although there was temporal variation from fish-to-fish, all swarms formed by: 1) the early recruitment of neutrophils to the inflammatory site (scouting), 2) the arrival of a pioneer neutrophil to the wound site (initiation), followed by 3) the directed migration of neutrophils towards the pioneer to form swarms (aggregation) (Figure 5, Supplemental Movie 6). Within minutes of injury, neutrophils began directed migration to the wound site (Figure 5A). This early scouting of neutrophils lasted on average 88 ± 24 minutes (data not shown) and is consistent with reports in zebrafish and mammalian systems which describe the recruitment of neutrophils close to the inflammatory site in response to chemoattractant gradients ^9,22^. Swarm initiation began when the pioneer neutrophil arrived at the wound site and changed behaviour during the scouting phase (Figure 5B), and ended when the first neutrophil joined the swarm (on average 36 ± 7 minutes, data not shown). During the aggregation phase, swarms developed by the directed migration of neutrophils, which lasted on average 183 ± 25 minutes, or until the end of the imaging period (Figure 5C). These stages are consistent between larvae and are comparable to the swarm stages reported in mammals ^7,9^. Taken together, these observations suggest that within the complexity of the inflamed tail fin, specific guidance cues are produced from a single pioneer neutrophil which promotes neutrophil swarming within the damaged tissue.

**Figure 5.**
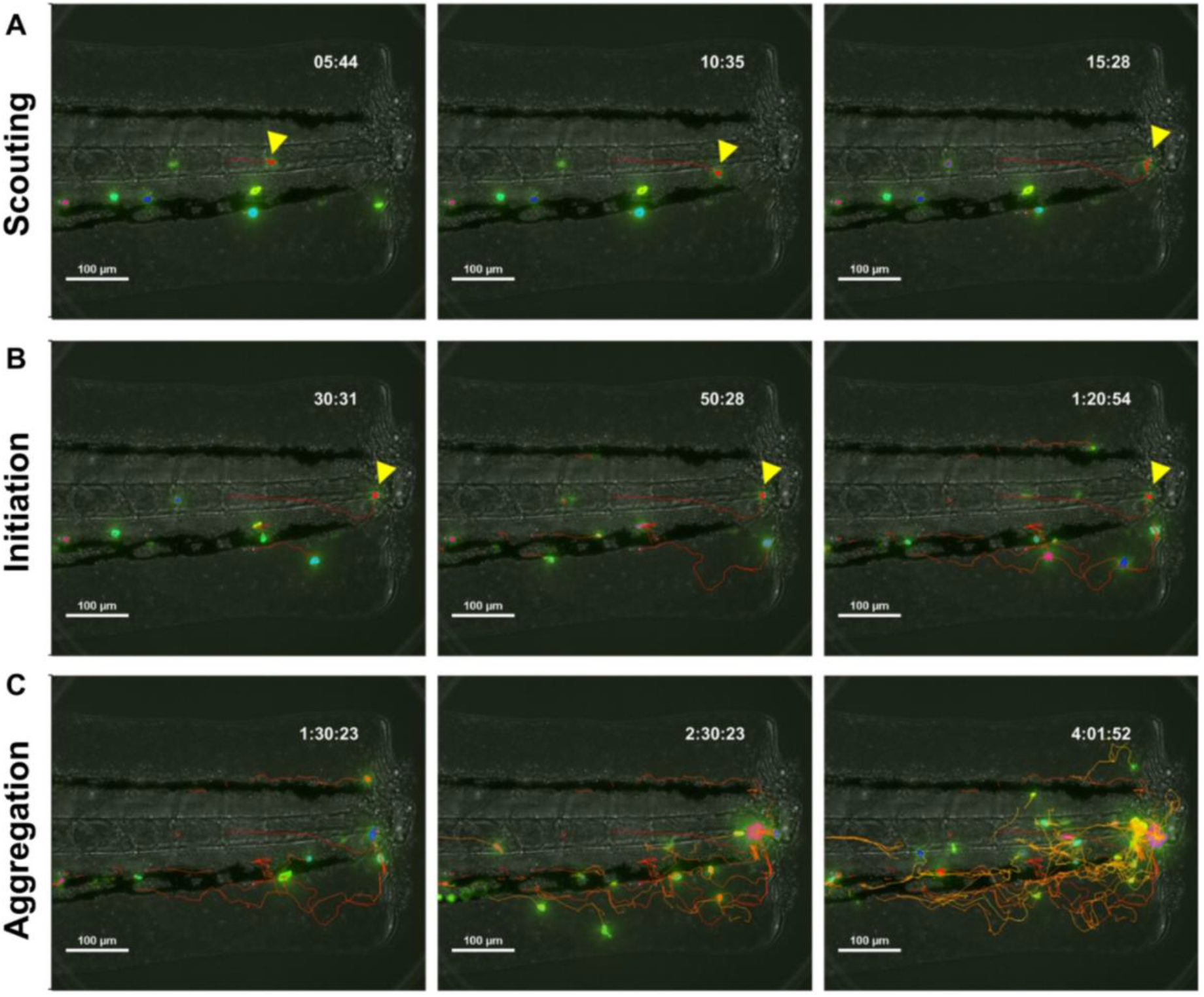
Neutrophil swarming responses to tissue damage occur in three sequential stages. Representative time-lapse sequence showing coordination of neutrophils to form swarms within the inflamed tail-fin. **A** Stage 1; scouting. The recruitment of neutrophils close to the wound site within minutes following tail fin transection. **B** Stage 2; initiation. Pioneer neutrophils migrate to the tissue region which becomes the swarm centre where they remain non motile. **C** Stage 3; aggregation. Swarming neutrophils direct their migration towards the pioneer neutrophil, resulting in swarm growth and neutrophil aggregation. Tracks are coloured by time where red corresponds to early and yellow corresponds to late arriving neutrophils. Time stamps are h:mm:ss relative to the start of imaging period at 30 minutes post injury.

### Pioneer neutrophils are viable prior to the onset of swarming

Cell death signals released from early-recruited pioneer neutrophils drive neutrophil swarming in mammals ^8,9,13^, although the precise signals and mode of cell death remain to be determined. We next used our zebrafish model to study cell death in pioneer neutrophils. DNA intercalating agent propidium iodide was used to determine pioneer neutrophil viability during the swarm initiation phase. Pioneer neutrophils excluded propidium iodide (Figure 6A-D), demonstrating that the plasma membrane of these cells remained intact prior to swarming and that these cells were viable. Interestingly, the tissue surrounding pioneer neutrophils was dense with extracellular DNA and cellular debris (Supplemental Movie 7). Apoptotic neutrophils would also exclude propidium iodide, therefore a FRET-based reporter for neutrophil apoptosis ^29^ was used to determine whether pioneer neutrophils were apoptotic. Analysis of pioneer neutrophils prior to swarming in *Tg(mpx:CFP-DEVD-YFP)sh237* larvae identified that despite the rounded, non-motile morphology, a FRET signal was present during both the scouting and initiation phases in all imaging runs where swarming was observed (Figure 6E, Supplemental Movie 8, n=6 neutrophils from 5 experimental repeats), demonstrating that pioneer neutrophils were not apoptotic. Neutrophil apoptosis at this early time point is rare, however when an apoptotic event was detected in neutrophils (n=2 independent observations), it was not followed by a neutrophil swarming response (Figure 6F, Supplemental Movie 9, n=2 neutrophils). These data demonstrate that pioneer neutrophil plasma membranes are intact prior to swarming, and that they are also not undergoing apoptosis. Together these findings suggest that pioneer neutrophils are viable prior to the onset of swarming.

**Figure 6.**
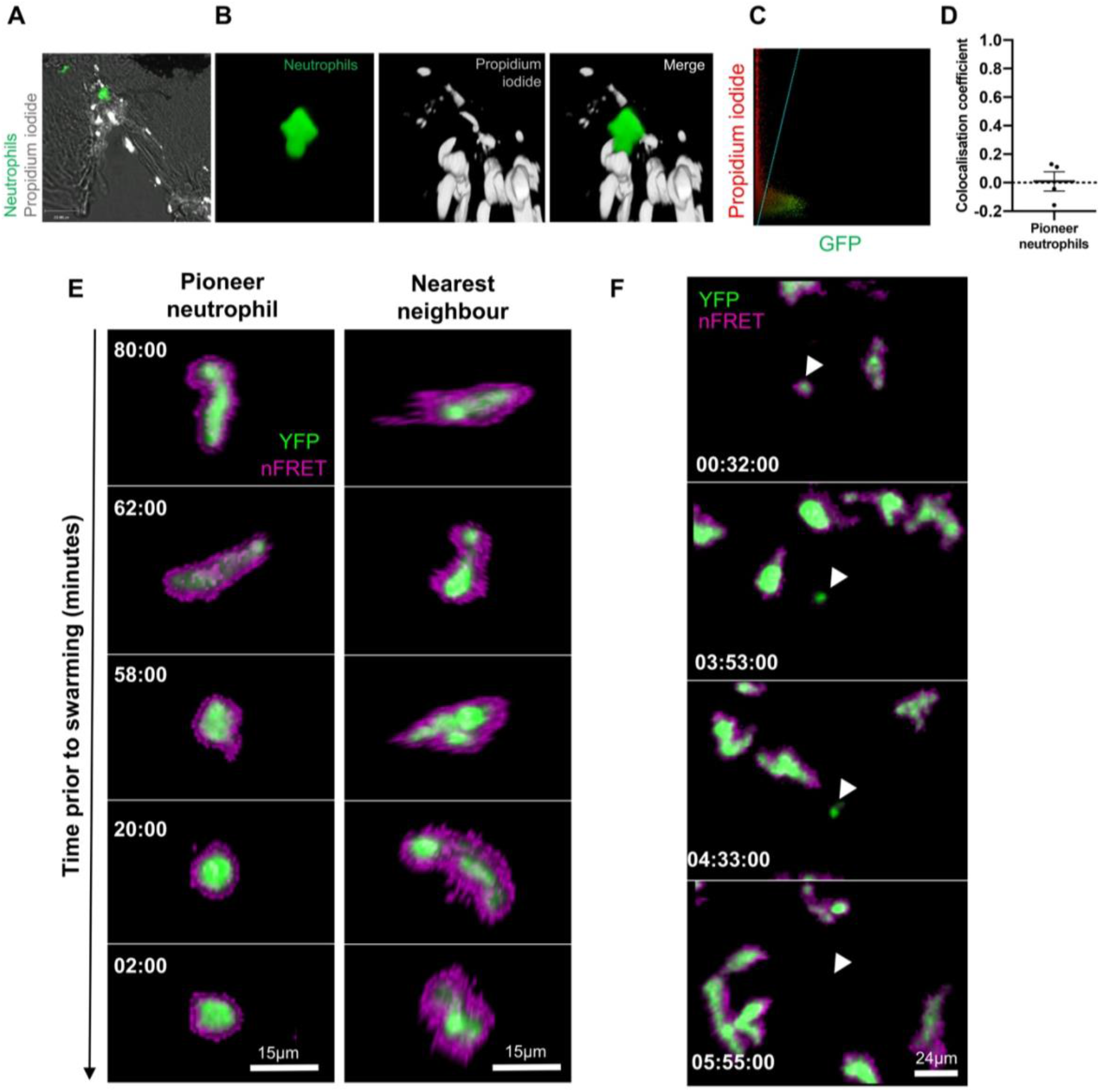
Pioneer neutrophils are viable prior to swarming. **A-D** Pioneer neutrophils are not propidium iodide positive prior to swarming. A Single slice image showing tail fin of injured *mpx*:GFP larva (bright field), stained with propidium iodide (grey). Image shows representative example of pioneer neutrophil at the wound site prior to the swarming response (green). **B** Representative 3D render of pioneer neutrophil during the initiation phase. Left tile shows *mpx*:GFP pioneer neutrophil, middle tile shows propidium iodide staining, right tile shows the two merged. **C** Representative colocalization analysis of pioneer neutrophils, where neutrophil signal (GFP) is on the x axis and propidium iodide signal (mCherry) is on the y axis. **D** Pearsons colocalisation coefficient for pioneer neutrophils (Data shown are mean ± SEM n=4 independent experiments). **E** Pioneer neutrophils are not apoptotic prior to swarming. 3dpf *mpx*:FRET larvae were injured and time lapse imaging was performed from 30 minutes post injury for 6 hours. Neutrophil signal from the acceptor (green) and nFRET (magenta) are shown to illustrate neutrophil apoptosis. Representative example of a pioneer neutrophil and its nearest neighbour in the frames preceding neutrophil swarming. The initiation stage is observed 58 minutes prior to swarming (rounded pioneer neutrophil). nFRET signal is in tact at all stages of migration prior to swarming in both the pioneer and nearest-neighbour non-pioneer neutrophil. Time stamps are mm:ss relative to the swarm start time. *(representative example of n=6 neutrophils from 5 independent repeats)*. **F** Apoptotic neutrophils do not initiate swarming. Example of neutrophil apoptosis at the wound site demonstrated by loss of FRET signal around 4 hours post injury, followed by the absence of neutrophil cluster formation in the same tissue region by the end of the imaging period. Time stamp is relative to injury time and is hh:mm:ss.

### Pioneer neutrophils release extracellular traps during swarming

Pioneer neutrophil cell death is accompanied by the appearance of cell fragments around aggregating neutrophils in mammals ^8,13^. During the aggregation phase of the swarming response in zebrafish, neutrophil fragments appeared around developing clusters (Figure 7A). To study these cell fragments, a high-resolution confocal microscopy approach was adopted (Supplemental figure 6). Strikingly, analysis of swarming neutrophils identified that large cytoplasmic vesicles, as well as smaller fragments of neutrophil debris, were released from swarms (Figure 7B-C, Supplemental Movie 10). This violent release of neutrophil fragments accompanied by large cytoplasmic structures, has been described associated with extracellular trap release in mammalian neutrophils ^30,31^. We therefore hypothesised that pioneer neutrophil cell death could be NETosis.

**Figure 7.**
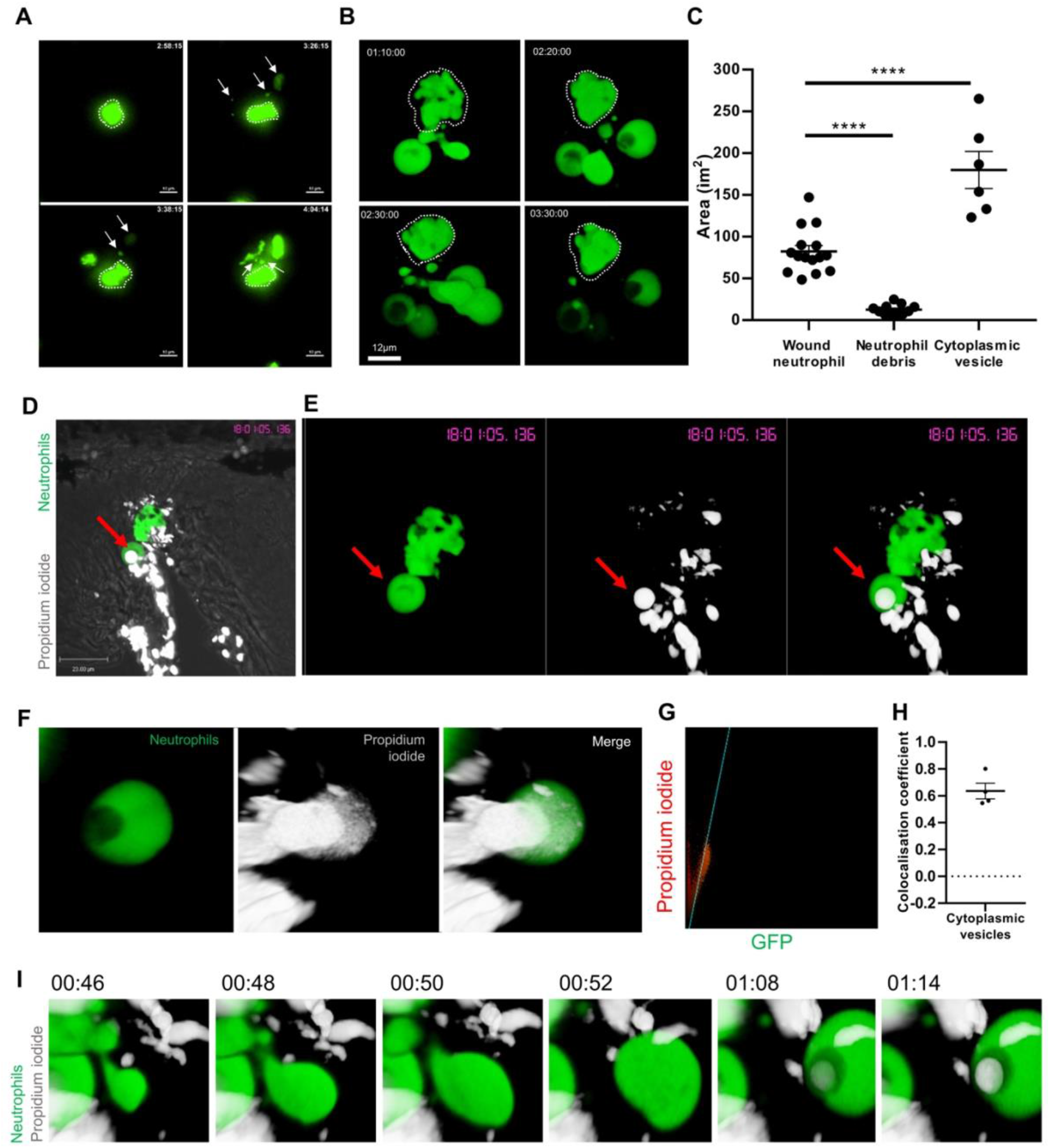
Catapult release of extracellular DNA by swarming neutrophils. **A** Maximum intensity projection time lapse sequence showing the production of cell fragments around swarming neutrophils (white arrows). Time stamps are h:mm:s relative to the start of the imaging period at 30 minutes post injury. Images were taken using a 20X objective on a Nikon widefield microscope using an andor zyla camera. **B** 3D rendered time lapse sequence showing cell fragments around swarming neutrophils (swarm highlighted by white dashed line). Time stamps are hh:mm:ss relative to injury time. Images were taken using a 40X objective lens on a perkin elmer spinning disk confocal microscope. **C.** The area of cellular debris and cytoplasmic vacuoles detected during swarm aggregation were measured alongside 3 nearest neighbour neutrophils at the wound site. (Error bars are SEM. Groups were analysed using an ordinary one way anova with Tukeys multiple comparison, p<0.0001. N=5 independent experiments). **D** Single z-slice of tail fin injury site labelled with propidium iodide (grey), illustrating neutrophil cytoplasmic vacuole (red arrow) released from a swarm. **E** 3D render showing propidium iodide positive neutrophil extracellular trap from swarming neutrophils. Images were taken using a 40X objective lens on a perkin elmer spinning disk confocal microscope. Time stamps are hh:mm:ss relative to injury time. **F-H** Colocalisation of propidium iodide with neutrophil cytoplasmic fragments. **F.** Representative image illustrating colocalization between the cell fragment and propidium iodide. **G** Representative Pearsons colocalization analysis of neutrophil extracellular traps, where GFP is on the x axis and mCherry is on the y. **H** Pearsons colocalisation coefficient for neutrophil extracellular traps released from swarming neutrophils (n=4). **I** Catapult-like release of extracellular DNA by individual neutrophil. Time course shows 3D render of a neutrophil at 00:46 stretching its cytoplasm resulting in the release of a large cytoplasmic vesicle which becomes positive for propidium iodide.

Neutrophil extracellular traps (NETs) are composed of a chromatin backbone embedded with antimicrobial proteins, which are able to capture and kill pathogens extracellularly ^17^. NETs were studied NETs in zebrafish using two approaches; neutrophil chromatin was labelled by making a transgenic zebrafish reporter for histone H2A under the neutrophil specific lyz promoter ^32^, and extracellular DNA was measured using DNA intercalating agent propidium iodide. High magnification confocal imaging was performed to study the cytoplasmic vesicle structures produced by swarming neutrophils. Propidium iodide labelled dead cells and extracellular DNA at sites of tissue injury (Figure 7D). Strikingly, the cytoplasmic structures produced alongside cellular debris became positive for propidium iodide, suggesting extracellular DNA is released by swarming neutrophils (Figure 7E-H, Supplemental Figure 8C, Supplemental Movie 11). These cytoplasmic structures were produced from an individual neutrophil following the stretching of neutrophil cytoplasm and violent release from neutrophils resulting in the production of small fragments of cellular debris and a large cytoplasmic structure which became propidium iodide positive (Figure 7I, Supplemental Figure 8A-B).

To confirm these structures contained histones, a transgenic reporter line for neutrophil histone H2A was generated to provide a cell-autonomous, intrinsically-expressed reporter of NET release *in vivo*. A genetic construct containing histone H2A with a C-terminal fusion of the fluorescent protein mCherry (H2A-mCherry), driven by the neutrophil specific *lyz* promoter ^32,33^ was generated using gateway cloning (Figure 8A). The construct was introduced into the genome of *mpx*:GFP larvae by Tol2 mediated transgenesis, and a stable was line generated: *TgBAC(mpx:GFP)i114;Tg(lyz:H2A-mCherry)sh530* (referred to as H2A-mCherry) (Figure 8B). The H2A transgene was expressed by neutrophils (Figure 8C), colocalising with the DNA stain DAPI within neutrophil nuclei (Figure 8D). The construct did not affect neutrophil migration to sites of inflammation (Figure 8E-F) demonstrating that the H2A-mCherry reporter was suitable for the study of neutrophil migration at sites of tissue injury. Analysis of swarming neutrophils in 3dpf H2A-mCherry larvae identified that NET-like structures were released by swarming neutrophils following the protrusion and stretching of the neutrophil cytoplasm, accompanied by the catapult-like release of histones from the nucleus resulting in the budding off of histone material from the releasing neutrophil (Figure 9). Together, these data demonstrate that the cell death associated with swarming is in part, NETosis, and provide novel insight into the moment of NET release *in vivo*.

**Figure 8.**
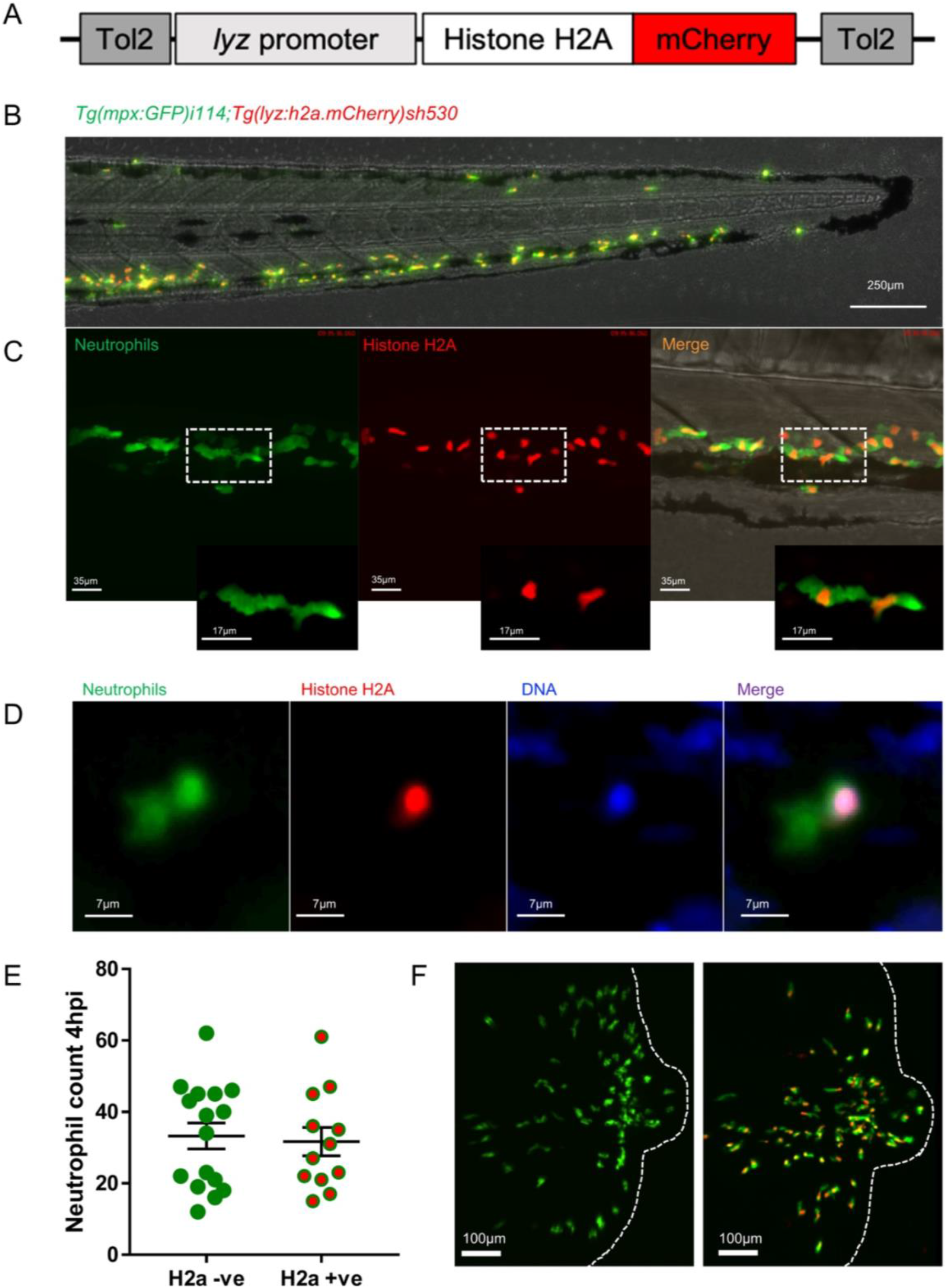
A transgenic zebrafish H2A.mCherry reporter for neutrophil extracellular traps. **A** Schematic of the lyz:H2A.mCherry construct made by Gateway cloning which includes the neutrophil specific promoter (lyz), and the histone H2A gene fused to the fluorescent protein mCherry flanked by Tol2 arms to aid transgenesis. **B-C** Representative image of the stable *TgBAC(mpx:GFP)i114;Tg(lyz:h2a.mCherry)sh530* transgenic line. **B** Image shows the caudal haematopoietic tissue of a 3dpf sh530 larvae, where the H2A mCherry transgene is expressed in neutrophils. **C** 40X confocal image of the transgenic line, showing neutrophil histones labelled by the transgene. **D** Dapi staining confirms that the H2A labels neutrophil DNA, image shows single z-slice of a CHT neutrophil from a 3dpf larvae. **E** Neutrophils counted at the wound site at the peak of recruitment (4 hours post injury, hpi) in H2A.mCherry negative and positive larvae. n=12-16 larvae from 3 independent experiments, p=0.77 where groups were analysed using an unpaired t-test. **F** Representative images of neutrophil recruitment to tissue injury (wound edge illustrated by white dashed line) in H2A.mCherry positive larvae and H2A.mCherry negative siblings. Images taken using a 20X spinning disk confocal microscope.

**Figure 9.**
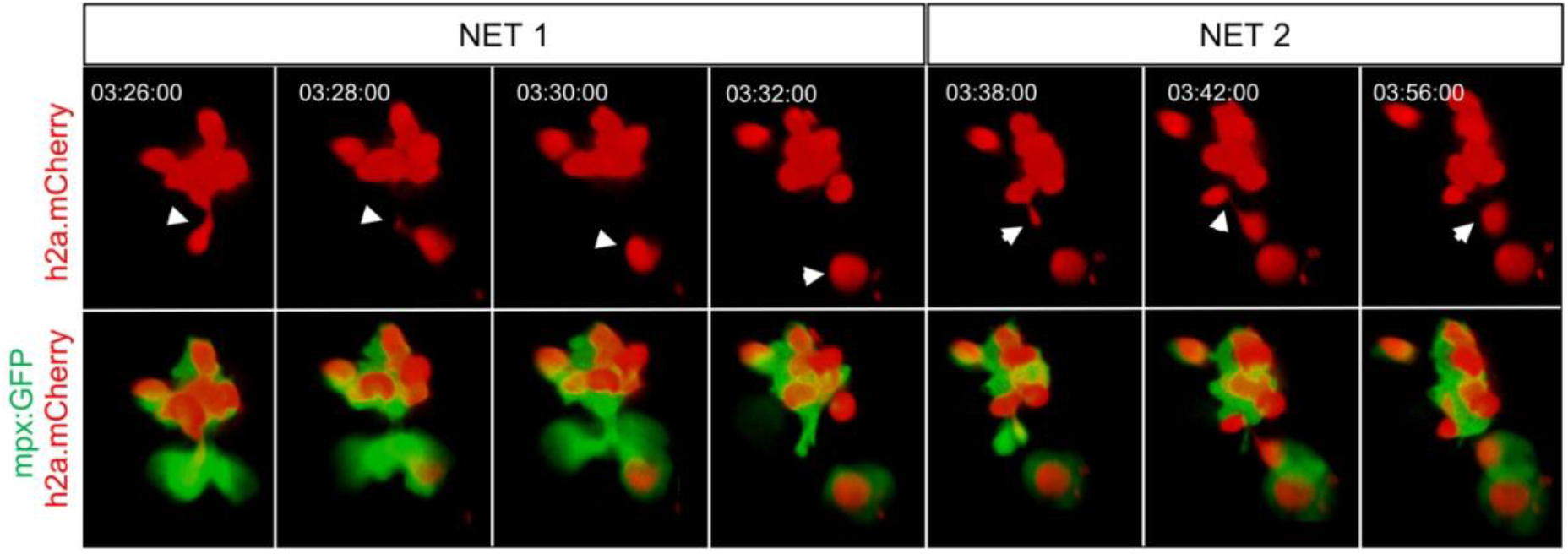
Histone H2A release from swarming neutrophils. Representative example of NET release from swarming neutrophils from 6 independent observations. Time course of *Tg(mpx:GFP)i114;Tg(lyz:h2a.mCherry)sh530* larva showing a single neutrophil and histone H2A (white arrows), undergoing NET-like morphological changes whereby histones are released from the centre of swarms in cytoplasmic vesicles. Time stamps are hh:mm:ss relative to time post injury.

After determining that NETs were released by early swarming neutrophils, we next determined whether pioneer neutrophils were amongst the NET releasing cells. The fate of pioneer neutrophils was studied within developing swarms using a photo conversion approach. Neutrophils were studied in a zebrafish reporter line which expresses the photoconvertible protein kaede specifically in neutrophils *Tg(UAS:Kaede);TgBAC(mpx:GAL4-VP16)i222* (referred to as *mpx*:kaede) ^19,34,35^. 3dpf *mpx*:kaede larvae were injured and the first neutrophil to the wound site was converted from green to red fluorescence. In larvae where photoconverted neutrophils became pioneers and swarms developed, pioneer neutrophils were studied within developing swarms (Figure 10A). Analysis of these neutrophils identified that photoconverted pioneer neutrophils underwent the morphological changes associated with NET release, with the production of a large cytoplasmic vesicles and neutrophil debris being observed from within the cluster (Figure 10B-C, Supplemental Movie 12). These data suggest that cell death by NETosis could be central to the swarming response, providing evidence of a role for NETosis in regulation of inflammation, aside from any direct role in host-defence.

**Figure 10.**
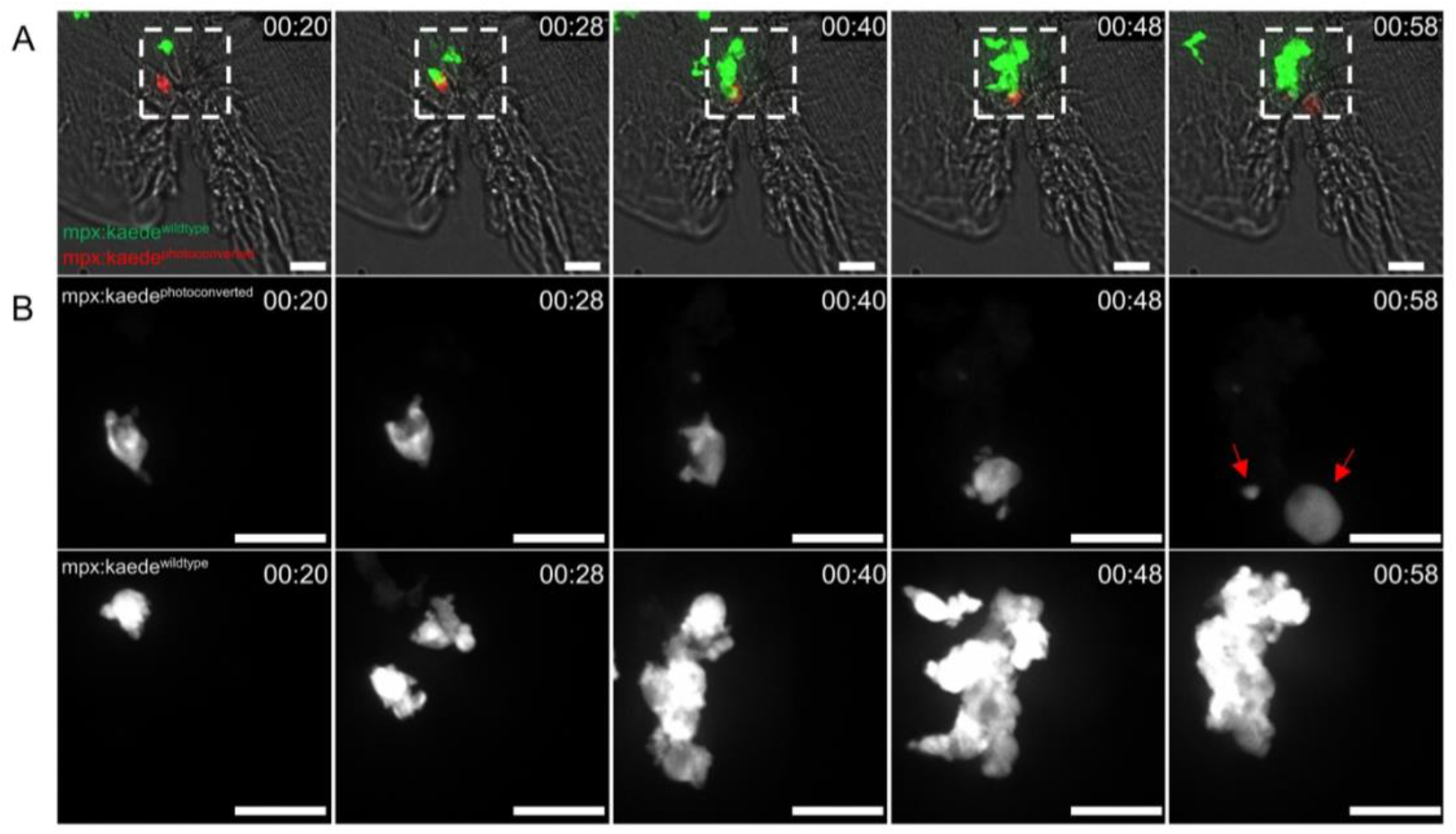
Pioneer neutrophils release NETs during the swarming response. **A** Photoconversion approach to study pioneer neutrophils within developing clusters. 3dpf *mpx*:kaede larvae were injured and the neutrophil closest to the wound site was photoconverted from green to red at 10 minutes post injury. Larvae where the red neutrophil was the swarm-initiating pioneer neutrophil were analysed. The large cytoplasmic structure and cellular debris associated with NETosis can be seen by 58 minutes post injury (red arrows). **B-C** Magnified region of interest highlighted by white dashed line. **B** Time course of photoconverted red neutrophil. **C** Time course of green wildtype swarming neutrophils which cluster around the red pioneer neutrophil. Images shown were taken using a 40X spinning disk confocal microscope. Scale bars are 12μm. Time stamps are hh.mm relative to time post injury.

## Discussion

In this study we investigated the migration patterns of neutrophils in the context of inflammation and infection and identified that neutrophil swarming behaviour is conserved in zebrafish immunity. We focused on neutrophil swarming in injury-induced inflammation, where the zebrafish model allowed us to track endogenous neutrophils in a physiologically-relevant tissue damage model *in vivo*. Utilising the optical transparency of zebrafish larvae and a combination of transgenic reporter lines and fluorescent DNA intercalating agents, we identified that swarm initiating pioneer neutrophils release extracellular traps from within swarms, building on a growing body of work that implicates a role for pioneer neutrophil death by NETosis in the swarming response ^8,9,13,36^.

We utilised the zebrafish model to precisely track neutrophils over time, providing some of the first *in vivo* characterisation of endogenous neutrophil migration patterns in the context of swarming at sites of tissue injury. Within the inflamed tail fin, we found that neutrophil swarms developed around an individual pioneer neutrophil which was sufficient to initiate swarming in zebrafish larvae. The pioneer neutrophils in our model share function with the pioneer neutrophils essential for swarm initiation in mice ^8,13,37^. Due to the relatively few number of neutrophils present in zebrafish larvae (~300) in comparison with the thousands (2–5×10^4^) ^9^ injected into the mouse ear, we propose that just one pioneer neutrophil is sufficient to drive a swarming response in our model.

The single-cell resolution achieved in our study enabled us to study pioneer neutrophils with optical clarity prior to the onset of swarming. Other groups have found that within inflamed or infected interstitial tissue, the initial arrest of a small number of ‘pioneer’ or ‘scouting’ neutrophils precedes a later influx of neutrophil migration ^8,10,13^ Based on our observations, we distinguished the pioneer neutrophil from other scouting neutrophils and propose that pioneer neutrophils have specialised functions required for swarm initiation, whilst scouting neutrophils are simply early responders to chemoattractants produced by damaged cells or pathogens at the inflammatory site.

Based on the morphology of pioneer neutrophils we investigated cell death signalling using transgenic reporter lines and cell viability dyes. Murine pioneer neutrophils are propidium iodide positive prior to the onset of swarming ^8,13^. Our data build on this observation, identifying that the propidium iodide signal comes from the cytoplasmic vesicle containing the neutrophil extracellular DNA. Interestingly, pioneer neutrophils themselves in our study were viable prior to swarming, as indicated by their exclusion of DNA intercalating agents, suggesting that lysis and release of cellular contents into the tissue is not an initiating factor in this model. We identify that pioneer neutrophils are surrounded by extracellular DNA, therefore it is tempting to speculate that initiating NETosis as the last effort of the neutrophil to capture and kill pathogens extracellularly would be an appropriate decision for a pioneer neutrophil. Perhaps the signals involved in initiating NETosis overlap with swarm initiation signals, whereby neutrophils call to other neutrophils for help as well as capturing pathogens by NETosis. Interestingly caspase-3 was intact during the swarm initiation phase, indicating that swarm initiating pioneer neutrophils were not undergoing neutrophil apoptosis prior to swarming. Due to the requirement for live imaging to study pioneer neutrophils prior to swarming, it was not technically possible to confirm our apoptosis results using staining assays such as TUNEL. However, other studies have found that results using the *mpx*:FRET transgenic line recapitulate TUNEL staining ^29^, demonstrating this is a reliable way to read out neutrophil apoptosis. These observations suggest that signals actively released from pioneer neutrophils initiate swarming, rather than the bursting of neutrophils and release of DAMPs into the tissue.

*In vivo* live imaging of NETs is achieved by labelling histones, neutrophils and extracellular DNA ^38^. Whilst imaging of NETs *in vivo* is advancing, our understanding of the kinetics of DNA release from neutrophils is still limited. There is growing evidence to suggest that NET release is conserved in zebrafish neutrophils, hence the zebrafish is a good model to investigate NET release ^39,40^. We developed an *in vivo* zebrafish reporter for NETs using histone H2A. Imaging data from other groups corroborate our identification that NET releasing neutrophils undergo distinct morphological changes involving the stretching of neutrophil cytoplasm and production of large extracellular DNA containing vesicles and cellular debris ^30,41^. In human neutrophils infected with *Staphylococcus aureus*, vesicles containing DNA are released into the extracellular space where they lyse and release their contents to form NETs, consistent with our observation that cytoplasmic vacuoles containing extracellular DNA are released by neutrophils ^41^. Furthermore, following LPS stimulation, murine neutrophils expel extracellular DNA in large cytoplasmic vesicles ^30^. We build on *in vitro* findings which suggests that DNA expelled from neutrophils diffuses around the releasing neutrophil, creating an extracellular trap with a large surface area ^31^. Furthermore, the catapult-like release of DNA and histones by neutrophils observed in our experiments is consistent with *in vitro* evidence that DNA release by eosinophils is catapult-like and expulsive ^42^. Our experiments enable the morphology of the NET releasing neutrophil to be observed *in vivo*, in real time, providing new insight into the morphological changes associated with NET release. These findings contribute *in vivo* data to the growing evidence that NETs facilitate neutrophil aggregation; neutrophil extracellular trap release is observed at sites of alum injection associated with neutrophil swarming in mice ^43^, and NET formation facilitates neutrophil aggregation at sites of fungal infection ^36^.

Neutrophil responses to tissue injury in murine systems are bi-phasic, and modulated in part by the lipid LTB4 which acts as a signal-relay molecule to amplify initial signals produced at inflammatory sites including formyl peptides ^44^. We demonstrate in our model that neutrophil recruitment to tail fin inflammation is bi-phasic; neutrophils proximal to the wound edge are recruited within minutes following injury, whilst neutrophils from further away recruited between 2-6 hours following injury. Using CRISPR/Cas9 to knock down *lta4h* and *blt1*, we found that neutrophil responses were impaired only in the later stages of recruitment (3-6hpi). These findings are in keeping with data from human and murine neutrophils ^7,8^. Neutrophil migration to form swarms can be further categorised into three phases: the early recruitment of neutrophils (referred to as ‘scouting’), followed by the large scale synchronised migration of neutrophils from distant regions (amplification), resulting in large scale tissue infiltration from the bloodstream (aggregation) ^7–9^. Consistent with these findings, neutrophil swarming at the wound site in our system occurred in three distinct stages, which are comparable to the sequential phases described in the swarming of neutrophils in intravenous/ intradermal transfer models in mice ^9,37^.

Our findings in this study implicate a role for pioneer neutrophil death by NETosis in swarm initiation. Understanding why swarms are initiated will be important for understanding the signals which control the coordination of neutrophil migration within interstitial tissue, which ultimately could lead to the identification of novel therapeutic targets for the treatment of chronic inflammatory disease.

## Materials and methods

### Zebrafish husbandry and ethics

To study neutrophils during inflammation *TgBAC(mpx:EGFP)i114, Tg(UAS:Kaede); TgBAC (mpx: GAL4-VP16) i222, Tg(mpx:CFP-DEVD-YFP)sh237*, and *TgBAC(mpx:GFP)i114;Tg(lyz:H2A-mCherry)sh530* zebrafish larvae were in-crossed. All zebrafish were raised in the Bateson Centre at the University of Sheffield in UK Home Office approved aquaria and maintained following standard protocols ^45^. Tanks were maintained at 28°C with a continuous re-circulating water supply and a daily light/dark cycle of 14/10 hours. All procedures were performed on embryos less than 5.2 dpf which were therefore outside of the Animals (Scientific Procedures) Act, to standards set by the UK Home Office.

### Tail fin transection assay

To induce an inflammatory response, zebrafish larvae at 2 or 3dpf were anaesthetised in Tricaine (0.168 mg/ml; Sigma-Aldrich) in E3 media and visualised under a dissecting microscope. For linear tail fin injury, tail fins were transected consistently using a scalpel blade (5mm depth, WPI) by slicing immediately posterior to the circulatory loop, ensuring the circulatory loop remained intact as previously described ^22^. For high resolution imaging, tail fins were nicked by placing the tip of the scalpel blade directly below the end of the caudal vein and slicing through the ventral fin, such that the entire wound site could be observed using a 40x objective.

### Widefield microscopy of transgenic larvae

For neutrophil tracking experiments, injured 3dpf *mpx*:GFP larvae were mounted in a 1% low melting point agarose solution (Sigma-Aldrich) containing 0.168 mg/ml tricaine immediately following tail fin transection. Agarose was covered with 500μl of a clear E3 solution containing 0.168 mg/ml tricaine to prevent dehydration. Time lapse imaging was performed from 0.5-5 hours post injury with acquisition every 30 seconds using 10 z-planes were captured per larvae over a focal range of 100μm using an Andor Zyla 5 camera (Nikon) and a GFP specific filter with excitation at 488nm. Maximum intensity projections were generated by NIS elements (Nikon) to visualise all 10 z-planes.

### Confocal microscopy of transgenic larvae

For visualising neutrophil swarming at high magnification, larvae were mounted in a 1% low melting point agarose solution (Sigma-Aldrich) containing 0.168 mg/ml tricaine for imaging immediately after tail fin transection. Agarose was covered with 2000μl of clear E3 solution containing 0.168 mg/ml tricaine to prevent dehydration. Imaging was performed from 30 minutes post injury using a 20x or 40x objective on an UltraVIEWVoX spinning disc confocal laser imaging system (Perkin Elmer), as indicated in figure legends. Fluorescence for GFP was acquired using an excitation wavelength of 488nm and emission was detected at 510nm, fluorescence for DAPI was acquired using an excitation wavelength of 405nm and emission was detected at 440nm and fluorescence for mCherry was acquired using 525nm emission and detected at 640nm. Images were processed using Volocity™ software.

### Tracking assays

Tracking of GFP labelled neutrophils was performed using NIS Elements (Version 4.3) with an additional NIS elements tracking module. A binary layer was added to maximum intensity projections to detect objects. Objects were smoothed, cleaned and separated to improve accuracy. A size restriction was applied where necessary to exclude small and large objects which did not correspond to individual neutrophils.

### Distance-time plots

For wound plots the distances from the wound were obtained by processing neutrophil tracks under the assumption that the tail fin wound is a straight line parallel to the x-axis of the greyscale image. Neutrophil tracking data was extracted from NIS elements and imported into MatLab software. For distance to pioneer plots the pioneer centre was set as a reference point and tracking was performed to determine neutrophil distance to the reference point. Tracks were extracted from NIS elements and plotted manually using GraphPad Prism version 7.0.

### Neutrophil specific expression of zebrafish genes

Gene expression was assessed using an RNA sequencing database from FACS sorted GFP positive cells from 5dpf zebrafish ^46^ (data deposited on GEO under accession number GSE78954). RPKM values for genes of interest were extracted. For single cell analysis gene expression values were extracted from the BASiCz (Blood atlas of single cells in zebrafish) cloud repository ^47^. Cells of the neutrophil lineage were analysed for expression of LTB4 signalling components.

### CRISPR/Cas9 reagents

Synthetic SygRNA® (crRNA and tracrRNA) (Merck) in combination with cas9 nuclease protein (Merck) was used for gene editing. Transactivating RNAs (tracrRNA) and gene specific CRISPR RNAs (crRNA) were resuspended to a concentration of 20μM in nuclease free water containing 10mM Tris-hcl ph8. SygRNA® complexes were assembled on ice immediately before use using a 1:1:1 ratio of crRNA:tracrRNA:Cas9 protein. Gene-specific crRNAs to target the ATG region of *blt1* and *lta4h* were designed using the online tool CHOPCHOP (http://chopchop.cbu.uib.no/). We used the following crRNA sequences targeting the ATG region of both genes, where the PAM site is indicated in brackets: *lta4h:* AGGGTCTGAAACTGGAGTCA(TGG), *blt1:* CAATGCCAATCTGATGGGAC(AGG).

### Microinjection of SygRNA^®^ into embryos

A 1nl drop of SygRNA^®^:Cas9 protein complex was injected into *mpx*:GFP embryos at the one cell stage. Embryos were collected at the one cell stage and injected using non-filament glass capillary needles (Kwik-Fil™ Borosilicate Glass Capillaries, World Precision Instruments (WPI), Herts, UK). RNA was prepared in sterile Eppendorf tubes. A graticule was used to measure 0.5nl droplet sizes to allow for consistency of injections. Injections were performed under a dissecting microscope attached to a microinjection rig (WPI) and a final volume of 1nl was injected.

### Genotyping and melting curve analysis

Site-specific mutations were detected using High Resolution Melting (HRM) Analysis which can reliably detect CRISPR/Cas9 induced indels in embryos ^48,49^. Genomic DNA extraction was performed on larvae at 2dpf. Larvae were placed individually in 0.2ml PCR tubes in 90μl 50mM NaOH and boiled at 95° for 20 minutes. 10μl Tris-HCL ph8 was added as a reaction buffer and mixed thoroughly. Gene specific primers were designed using the Primer 3 web tool (http://primer3.ut.ee/). Sequences were as follows *lta4h_fw:* CGTGTAGGTTAAAATCCATTCGCA *lta4h_rev:* GAGAGCGAGGAGAAGGAGCT *blt1_fw:* GTCTTCTCTGGACCACCTGC *blt1_rev:* ACACAAAAGCGATAACCAGGA. HRM analysis (Bio-Rad) PCR reactions were made with 5μl Sybr™ Green master mix (Thermo Fisher), 0.5μl of each primer (10μM), 1μl gDNA and 3μl water to make a final reaction volume of 10μl. PCR reactions were performed in a LightCycler instrument (Bio-Rad) using 96-well plates. The two-step reaction protocol was as follows: 95 °C for 2 min, followed by 35 cycles of 95 °C for 10 seconds, 58° for 30 seconds, 72° for 20 seconds. The second stage of the protocol was 95 °C for 30 seconds, 60 °C for 60 seconds, 65 °C for 10 seconds. The temperature then increased by 0.02 °C/s until 95 °C for 10 seconds. Melt curves were analysed using Bio-Rad software version 1.2. Succesful detection of CRISPR/Cas9 induced indels is illustrated in Supplemental figure 6. Mutagenesis frequencies of 91% and 88% were detected for *lta4h* and *blt1* respectively.

### *Staphylococcus aureus* preparation

*Staphylococcus aureus* strain SH1000 pMV158mCherry was used for all experiments^50^. An overnight bacterial culture was prepared by growing 1cfu of SH1000 pMV158mCherry in 10mLs of bovine heart medium (BHI) (Sigma Aldrich lot number 53286) and 10μLs of 5mg/mL tetracycline (Sigma-Aldrich) for 16-18 hours at 37°C. 500μLs of this overnight culture was then aliquoted into 50mLs of BHI (Sigma Aldrich, 53286) infused with 50μLs of 5mg/mL tetracycline (Sigma Aldrich) and grown until an optical density at 600nm of 0.5 was obtained. This culture was pelleted and resuspended in PBS (pH 7.4) (Fisher Scientific lot number 1282 1680) to a concentration of 2500cfu per nL.

### Otic vesicle injection

2500cfu of Sh1000 pMV158mCherry was injected into the left otic vesicle of 2dpf *Tg(mpx:GFP)i114* larvae. Injections were performed under a dissecting microscope attached to a microinjection rig (WPI) and a final volume of 1nl was injected. For analysis of swarm volumes larvae were fixed in 4% paraformaldehyde in PBS and imaged using a spinning disk confocal microscope.

### Staining and colocalization experiments

To study the plasma membrane integrity of neutrophils, 3dpf *mpx*:GFP larvae were incubated in 1% LMP agarose solution containing 0.1% propidium iodide (Sigma-aldrich) immediately following tail fin transection. Pearsons colocalisation analysis was performed by drawing a region of interest around neutrophil cytoplasmic vesicles using Volocity™ software. For DAPI staining of sh530 larvae, 2dpf larvae were fixed in 1ml of 4% paraformaldehyde (PFA) at room temperature for 30 minutes, washed in PBST and transferred to 100% MeOH overnight at −20. Samples were washed in PBST twice before permeabilization using proteinase K (10 μg/ml) for 20 minutes at room temperature. Samples were fixed for 20 minutes in 4% PFA at room temperature and washed twice in PBST. Samples were stained in a 0.1% DAPI (Sigma Aldrich) solution in 1X PBS for 20 minutes and kept in the dark. Samples were washed in PBST and imaging was performed.

### Förster resonance energy transfer imaging of neutrophil apoptosis

Neutrophil apoptosis was studied using our transgenic *Tg(mpx:CFP-DEVD-YFP)sh237* ^29^ zebrafish line which expresses a genetically encoded Förster resonance energy transfer (FRET) biosensor consisting of a caspase-3 cleavable DEVD sequence flanked by a CFP YFP pair^51^, under the neutrophil specific *mpx* promoter. A loss of FRET signal in this system provides a read out of apoptosis specifically in neutrophils *in vivo* in real time. To visualise apoptotic events in the context of neutrophil swarming, 3dpf *Tg(mpx:CFP-DEVD-YFP)sh237* larvae were injured and mounted in a 1% agarose solution containing 0.168 mg/ml tricaine and covered with 500μl of a clear E3 solution containing tricaine to prevent dehydration. FRET imaging was performed from 30 minutes post injury for 5 hours using a 20x objective lens on an UltraVIEWVoX spinning disc confocal laser imaging system (Perkin Elmer) with acquisition every 2 minutes. 10 z-planes were captured per larvae over a focal range of 100μm using the following filters: a donor CFP channel (440nm for excitation, 485nm for detection), an acceptor YFP channel (514nm for excitation and 587nm for detection), and a FRET channel (440nm for excitation and 587nm for detection). An Ultraview dichroic mirror passes 405,440,515,640 was used to increase imaging speed using these filter blocks. Volocity™ software was used to calculate normalised FRET values (nFRET). To compensate for the bleed through of the CFP and YFP fluorophores into the FRET channel, FRET bleed through constants were calculated. Control samples containing HeLa cells transfected with CFP alone or YFP alone were imaged using the same settings used for data acquisition of the *mpx*:FRET zebrafish reporter line. ROIs were drawn around a population of cells in the frame and Volocity™ software calculated FRET bleed through values as the mean intensity of the recipient channel (FRET) divided by the mean intensity of the source (CFP or YFP). These FRET constants were then used by Volocity™ to calculate a normalised FRET value. Neutrophil apoptosis was observed by overlaying the YFP and nFRET channels.

### Generation of histone H2A transgenic reporter line

A genetic construct containing histone H2A with a C-terminal fusion of the fluorescent protein mCherry, driven by the neutrophil specific *lyz* promoter was generated using gateway cloning ^52^. The final construct (pDestTol2CG2 lyz:H2A-mCherry cmlc2:EGFP) was assembled from the following gateway vectors: a 5’ vector p5E-MCS lyz containing 6.6kb of the lysozyme C promoter ^32^, a middle entry vector pME-H2AmCherry containing zebrafish histone H2A fused to mCherry, and a 3’ vector containing a polyadenylation site p3E-polyA. The final construct containing the Tol2 arms and green heart marker for easy recognition of successful transgenesis was created by an LR reaction combining the three vectors with the destination vector pDestTol2CG2. The construct was injected into transgenic *TgBAC(mpx:GFP)i114* larvae using Tol2 mediated transgenesis and a stable f2 line *TgBAC(mpx:GFP)i114;Tg(lyz:H2A-mCherry)sh530* was generated.

### Photoconversion of pioneer neutrophils

Photoconversion assays were performed using larvae expressing the photoconvertible protein kaede under the neutrophil specific *mpx* promoter: *TgBAC(mpx:GAL4-VP16); Tg(UAS:Kaede)i222* ^34^. At 3dpf larvae were anaesthetised and injured using the minor tail fin nick, and mounted immediately in a 1% LMP agarose solution containing tricaine. At ten minutes post injury, a region of interest was drawn around the neutrophil nearest to the injury site for photoconversion from green to red fluorescence. Photoconverstion of kaede labelled neutrophils at the wound site was performed using an UltraVIEWPhotoKinesis™ device (Perkin Elmer and Analytical Sciences) on an UltraVIEWVoX spinning disc confocal laser imaging system (Perkin Elmer). The photokinesis device was calibrated using a coverslip covered in photobleachable substrate (Stabilo Boss™, Berks UK). Photoconverstion was perfomed using a 405nm laser at 40% using 120 cycles, 250 pk cyles and 100ms as previously published ^34^. Successful photoconversion was detected through loss of emission detected following excitation at 488nm, and gain of emission following 561nm excitation. Following photoconverstion, timelapse imaging was performed from 20 minutes post injury for 4 hours. Photoconverted neutrophils which became swarm initiating pioneer neutrophils were analysed.

### Statistical analysis

Data were analysed using GraphPad Prism version 7.0. Paired *t* tests were used for comparisons between two groups and one-way ANOVA with appropriate post-test adjustment was used for comparisons of three or more groups.

## Acknowledgements

The authors would like to thank The Bateson Centre Aquarium Team at the University of Sheffield for their assistance with zebrafish husbandry. Imaging work was performed at the Wolfson Light Microscopy Facility, microscopy studies were supported by an MRC grant (G0700091) and a Wellcome Trust grant (GR077544AIA). We are extremely grateful to Dr. Tomasz Prajsnar for providing *S. aureus* strains.

## Competing Interests

The authors declare no conflict of interest.

## Funding Information

This work was supported by a Medical Research Council (MRC) Senior Clinical Fellowship with Fellowship-Partnership Award and MRC Programme Grants to S.A.R (G0701932 and MR/M004864/1) and an MRC Centre Grant (G0700091). P.M.E is funded by a Sir Henry Dale Fellowship jointly funded by the Wellcome Trust and the Royal Society (Grant Number 105570/Z/14/Z).

---

**Supplemental Figure 1.**
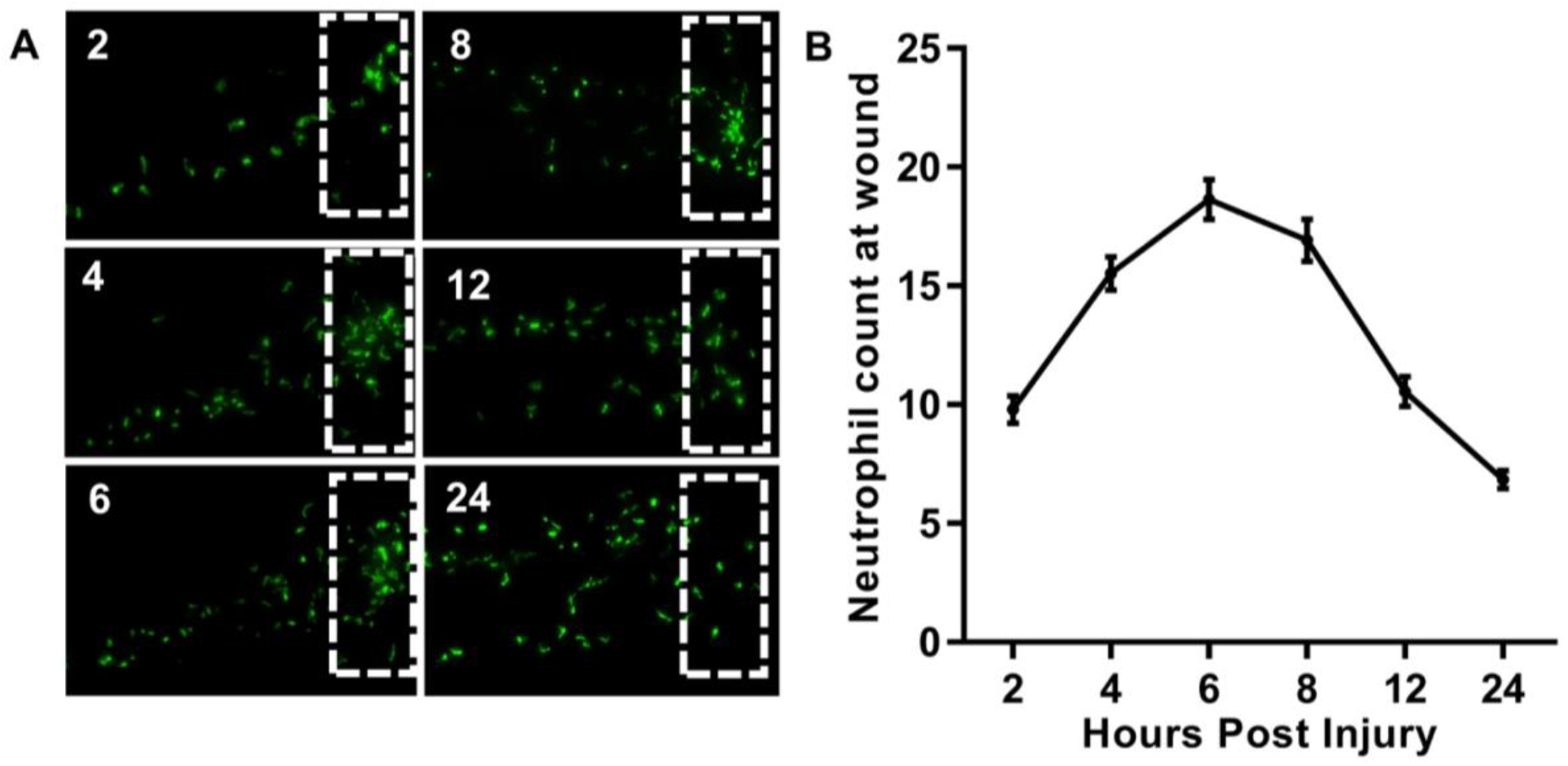
Dynamics of the neutrophil response to tissue injury. Following tail-fin transection of *mpx*:GFP transgenic larvae, the number of GFP neutrophils at the site of injury were counted at 2, 4, 6, 8,12 and 24 hours post injury. **A** Representative images illustrating neutrophils in the tail fin region throughout the inflammatory time course. **B** Quantification of neutrophil counts at the wound site throughout the time course. Data are shown as mean ± SEM, n= 53 larvae from 3 experimental repeats.

**Supplemental Figure 2.**
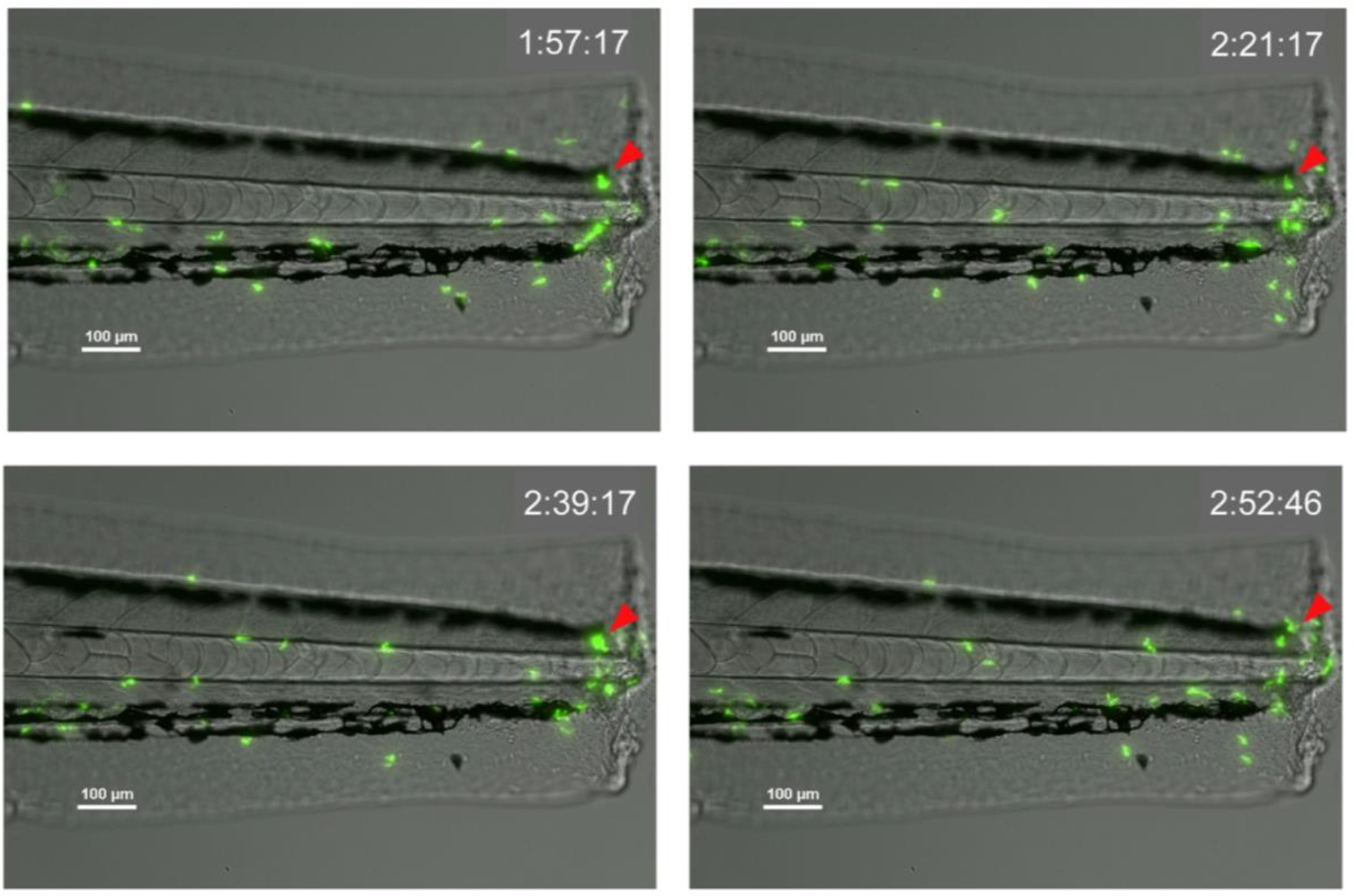
Transient neutrophil swarms are observed within the inflamed tail fin. Time course of *mpx*:GFP transgenic zebrafish larvae following tail-fin transection illustrating short-lived (<1 hour) transient neutrophil swarming at the wound site. Phases of coordinated migration resulting in cluster formation (red arrow) were observed within the imaging period, followed by cluster dissipation and re-formation. Time stamps shown in white (h:mm:ss) are relative to the start of imaging at 30 minutes post injury.

**Supplemental Figure 3.**
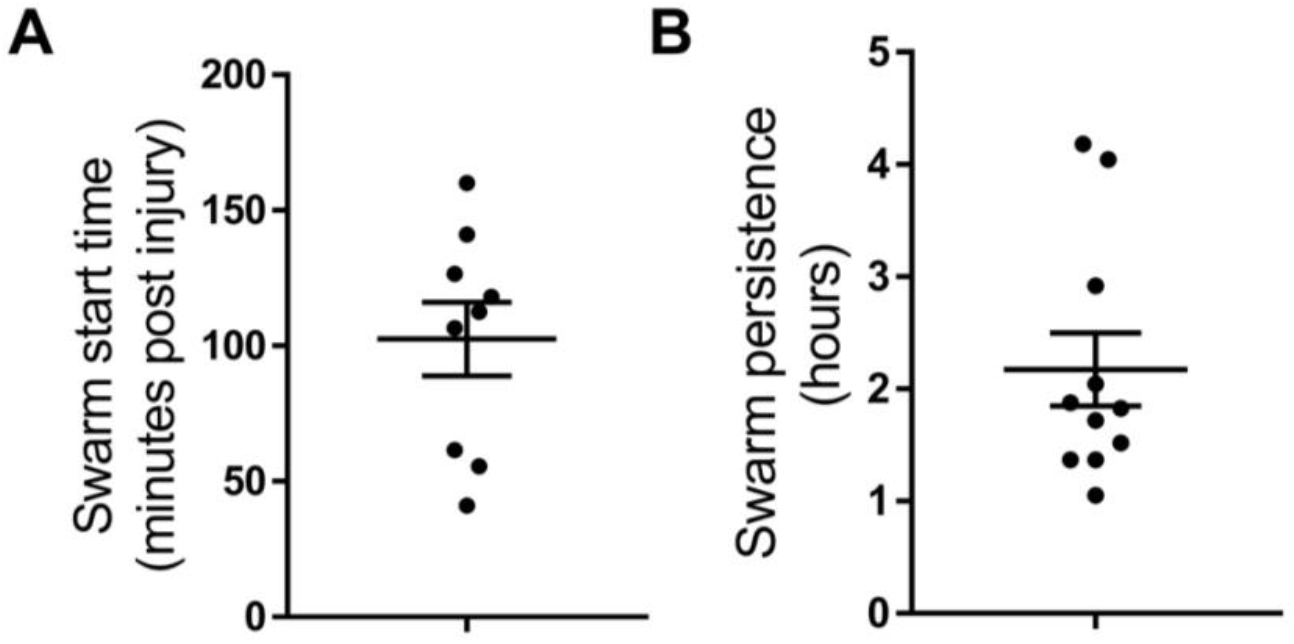
Characterisation of persistent neutrophil swarms. **A** Time (minutes post injury) in which persistent neutrophil swarms began to develop following tail-fin transection in zebrafish larvae (n=5 experimental repeats). **B** Persistence time of neutrophil swarms measured during 5 hour imaging period (n=5 experimental repeats).

**Supplemental Figure 4.**
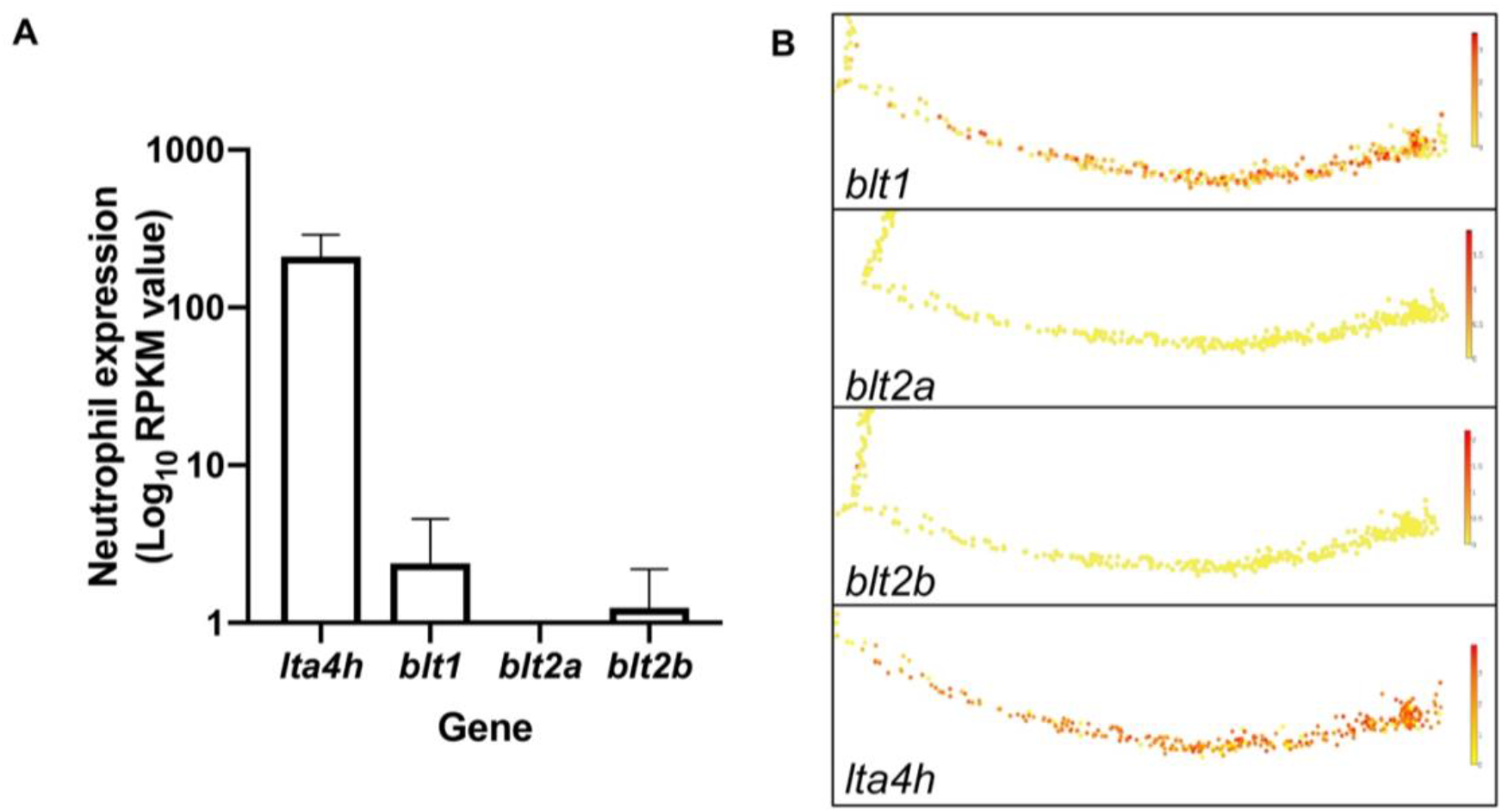
Expression of LTB4 signalling components in zebrafish neutrophils. **A** RNA sequencing of FACS sorted GFP positive cells from 5dpf *mpx*:GFP zebrafish larvae. RPKM values illustrate zebrafish neutrophil expression of *lta4h, blt1, blt2a* and *blt2b*. **B** Single-cell gene expression profiles of LTB4 signalling components expressed in the zebrafish neutrophil lineage, extracted from the Sanger BASiCz zebrafish blood atlas. Circles represent individual cells colour coded where red is high expression and yellow is no expression.

**Supplemental Figure 5.**
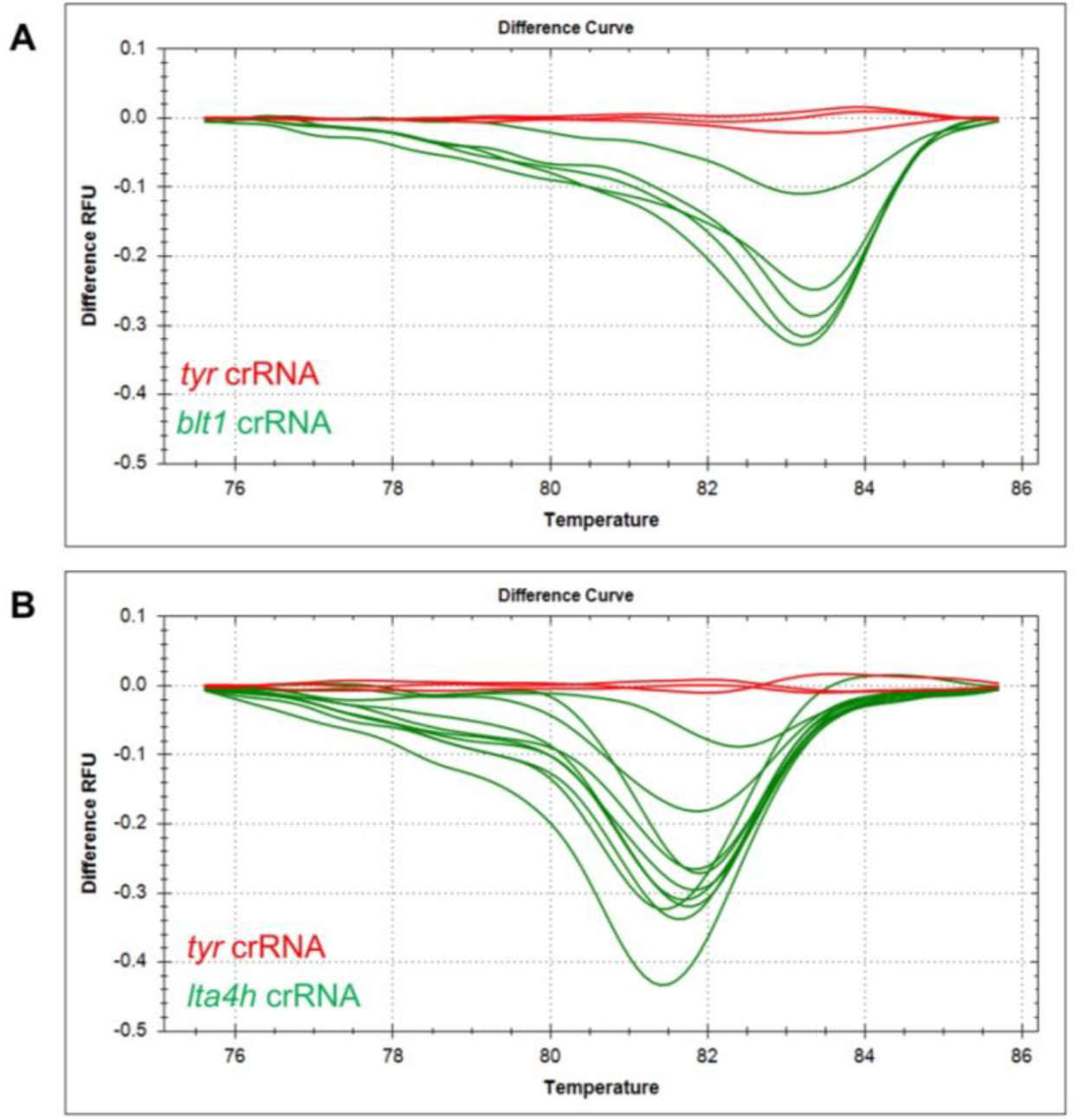
High resolution melt curve analysis for genotyping *blt1* and *lta4h* Crispants. Genotyping example of successful CRISPR-induced indels by high resolution melt analysis for *blt1* (**A**) and *lta4h* (**B**) injected larvae. Wild type curves (red) from three representative control *tyrosinase* larvae and shifted, irregular melt curves (green) corresponding to mosaic heteroduplex PCR fragents formed as a result of CRISPR/Cas9 mutations.

**Supplemental Figure 6.**
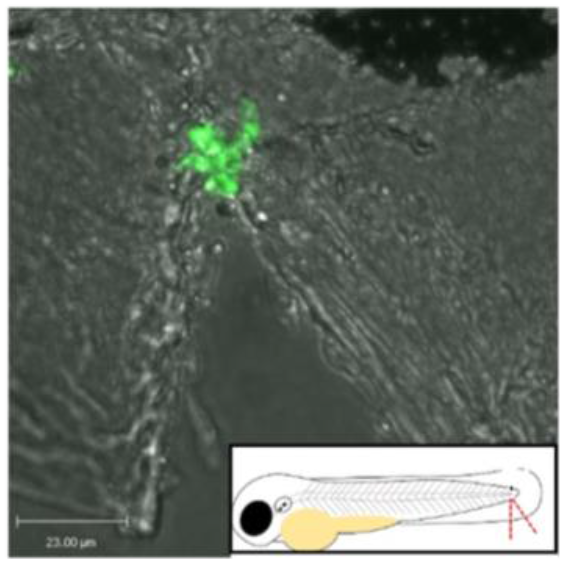
High magnification confocal microscopy tail fin injury model. Tail fin nick was performed on 3dpf *mpx*:GFP larvae which were imaged using a 40X objective lense on a perkin elmer spinning disk confocal microscope.

**Supplemental figure 8.**
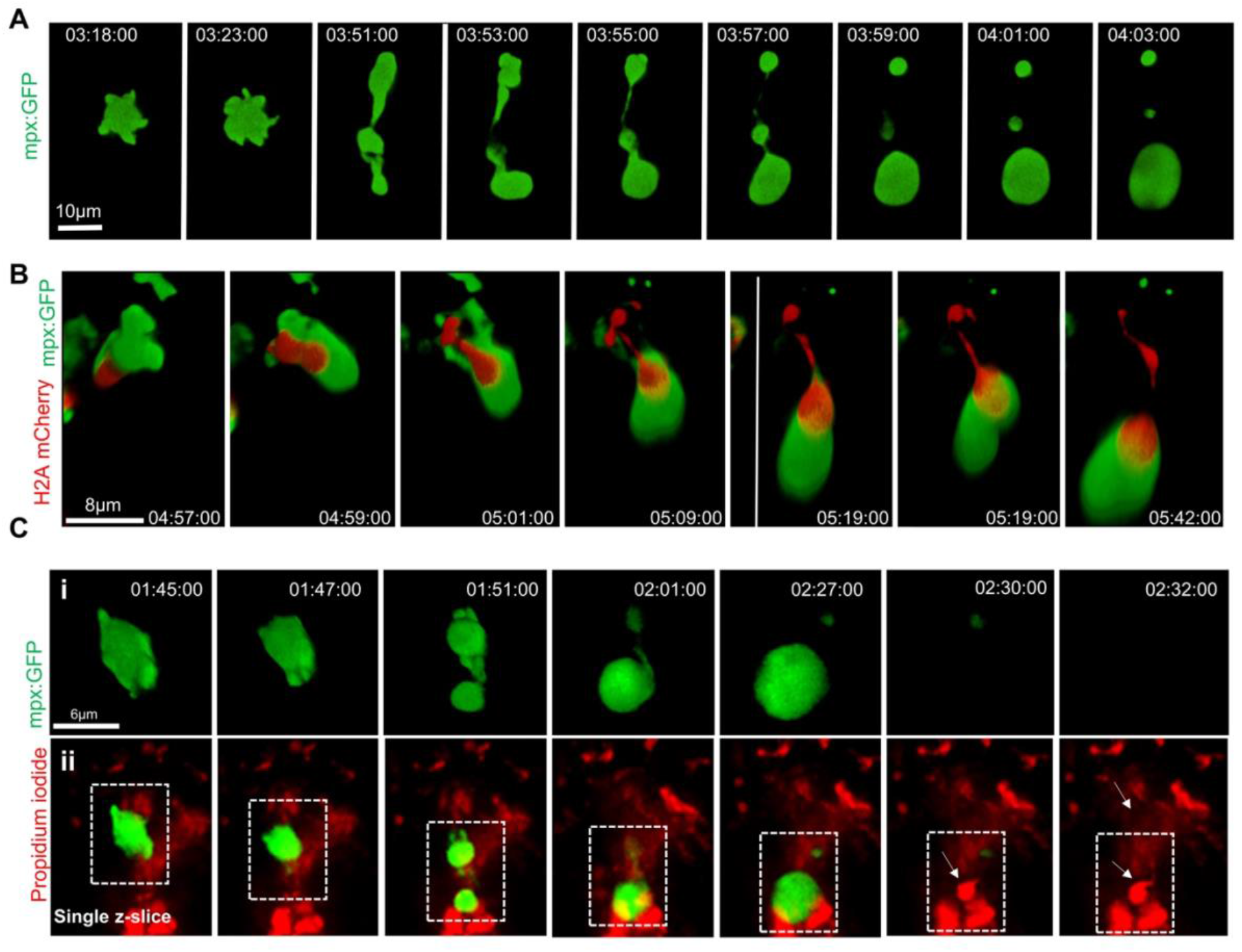
Dynamic imaging of the moment of NET release. **A** Time course of a neutrophil in a *TgBAC(mpx:GFP)i114* transgenic larva showing a single neutrophil undergoing NET-like morphological changes at the wound site, as illustrated by the production of cloud-like vesicle and cellular debris. **B** Time course of *Tg(mpx:GFP)i114;Tg(lyz:h2a.mCherry)sh530* larva showing a single neutrophil and histone H2a, undergoing NET-like morphological changes. **C** Time course of *TgBAC(mpx:GFP)i114* transgenic larva stained with propidium iodide (red). **i.** NET-like cell death accompanied by the release of cloud-like vesicle and cellular debris. **ii.** Single z-slice illustrating the cloud-like vesicle becoming propidium iodide positive, demonstrating extracellular DNA release. Unless stated, data shown are 3D reconstructions of image sequences taken using a 40X objective lens on a spinning disk confocal microscope. Time stamps are hh:mm:ss relative to time post injury.

## Notes

#### Summary of Updates

Updated extended version including work on neutrophil extracellular traps.

